# A *Mycobacterium tuberculosis* secreted virulence factor disrupts host snRNP biogenesis

**DOI:** 10.1101/2024.02.06.579120

**Authors:** Komal Chauhan, Nishat Passricha, Yogita Kapoor, Dipanwita Datta, Naresh Arora, Mrinmoy Das, Kavya Rao, Lakshyaveer Singh, Ravi Dutt, Raman Deep Sharma, Binayak Sarkar, Basanti Malakkar, Haroon Kalam, Prince Saini, Lalita Mehra, Prasenjit Das, Vineet Ahuja, Amit Singhal, Vinay Nandicoori, Dhiraj Kumar

## Abstract

We earlier reported that *Mycobacterium tuberculosis* (*Mtb*) alters host RNA splicing to help its intracellular growth. Here, we report an unprecedented mechanism whereby a secreted virulence factor from *Mtb* interferes with the biogenesis of key spliceosomal components, causing an altered RNA splicing pattern. A high-throughput yeast-2-hybrid screen identified several *Mtb*-secreted proteins that can interact with the host RNA splicing factors (SFs). Through custom-designed in-cell assays, we show that one of those proteins, Rv1435c, targets specific exon-skipping events to alter RNA splicing. We show that Rv14345c or **h**ost **s**plicing **r**egulator 1 (hsr1) facilitates direct interaction between *Mtb* phagosomes and U5 snRNA and SNRPF, key components of the snRNPs. Genetic deletion of *hsr1* reverses the specific exon-skipping events caused by WT *Mtb* infection. The *Δhsr1* strain shows compromised growth during *ex vivo* infection in the macrophages and *in vivo* infection in the mice. Tissue sections from the WT *Mtb* or *Δhsr1*-infected mice showed significant hsr1-dependent SNRPF staining, a phenomenon also noted in the human intestinal tuberculosis (ITB) biopsies. We infer that hsr1 is a virulence factor which alters RNA splicing by interacting with U5snRNA and SNRPF. The splicing regulators from the host and pathogen are novel targets for anti-tuberculosis therapy.

## Introduction

Tuberculosis (TB) caused by the bacterial pathogen *Mycobacterium tuberculosis* (*Mtb*) is a considerable challenge to the global sustainable development goals on health. With more than 10 million new TB cases and nearly 1 million deaths globally, the conventional approaches for prevention and control are increasingly looking insufficient to eradicate TB. The emerging paradigm on novel TB therapy envisages exploiting host-directed therapy (HDT). The advantage of HDT is its perceived universality in targeting drug-resistant or drug-sensitive infections. Also, HDTs, combined with antibiotics, are expected to hinder the development of new drug-resistant strains. Over the past two decades, several studies identified and reported host-dependency factors that help the intracellular growth and survival of *Mtb*, mostly by modulating the anti-bacterial functions of the hosts ^1–7^. In a genome-wide analysis of identifying such host factors, proteins from diverse functional classes, including gene expression, were reported to have pro-bacterial functions ^6^.

The transcriptional response of the host cells to *Mtb* infections has been studied extensively^8–12^. While the immediate response of host cells following infection with *Mtb* involves processes like trafficking, signalling and metabolic alterations, the long-term shaping of host response is decided by reshaping the transcriptional landscape of the cells^13,14^. Eukaryotic gene expression involves several post-transcriptional regulatory events, including capping, poly-adenylation, RNA splicing, etc. We previously showed that in the *Mtb*-infected macrophages, several genes classically known for their anti-bacterial functions were spliced in a manner that did not allow their translation or the polypeptides thus formed were truncated^13^. Moreover, the alternatively spliced transcripts often lacked critical functional domains due to exon exclusion events^14,15^. Several other studies together converge to the notion that RNA splicing could be a central vulnerability for the pathogens to exploit.

However, the molecular mechanism that allows pathogens, including *Mtb*, to alter host RNA splicing in a targeted manner remains poorly understood. RNA splicing is an evolutionarily conserved mechanism among eukaryotes and involves the assembly of a large multi-molecular complex called spliceosome at the exon-intro junctions^16–22^. The key components of the spliceosomes are small nuclear ribonuclear proteins or snRNPs, comprising snRNA and associated proteins^23–31^. Interestingly, snRNPs are formed through complex protein-RNA interactions spanning across the nucleus and the cytosol^29,32^. In addition, accessory proteins like hnRNPs, SR proteins and SRSF proteins play an important role in recognizing the exon-intron junctions and splicing reactions^17,19,33,34^. Cumulatively, there are plenty of potential steps where secreted virulence proteins from *Mtb* could intercept the RNA splicing machinery, thereby altering the course of downstream reactions. Secreted virulence factors of *Mtb* are classically known to interfere with cytosolic events like trafficking, antigen presentation, bioenergetics and nuclear events like epigenetic regulations ^35–39^.

To understand how *Mtb* alters host RNA splicing, we hypothesized that certain secreted virulence factors of *Mtb* could interact with and alter the assembly and/or recognition of spliceosome complexes or their components. Here, through a high throughput yeast-2-hybrid screening between secreted *Mtb* proteins and host splicing-related proteins or splicing factors (SFs), we identify several interacting partners between the two. We rationally narrowed it down to one *Mtb* protein, Rv1435c, to understand the molecular mechanism in detail. Using a genetic knockout of Rv1435c, we show that cytosolic interactions with snRNP biogenesis components allowed *Mtb* to alter the stoichiometry of snRNPs in the cells, eventually altering the RNA splicing process.

## Results

### Dysregulated spliceosome complex composition in *Mtb*-infected macrophages

Previously, we reported a targeted alteration of host RNA splicing pattern upon *Mtb* infection in the macrophages^13^. We also showed how altered RNA splicing impacted cellular function since a lot of target genes were spliced in a manner that led to the mRNAs, which were either non-functional (lacking specific functional domains), unstable or could not get translated (Fig. 1A)^14^. These alterations were reported as advantageous for the pathogen inside the host macrophages. To understand how RNA splicing was specifically altered in the infected macrophages, we systematically analyzed the RNA splicing process in these cells. RNA splicing requires the assembly of a large multimolecular complex, called the spliceosome, in a step-wise manner, which helps in the recognition, excision of the introns and stitching of the exons together (Fig. 1B). To analyze the assembly of spliceosome complexes in the *Mtb* infected macrophages, we decided to purify and analyze the *in vivo* assembled spliceosome complexes from these cells. THP-1 macrophages were infected with H37Rv (the laboratory strain of *Mtb*) at 1:10 MOI, and at 48 hours post-infection (h.p.i.), cytosolic and nuclear extracts were prepared without disrupting the assembled spliceosome complexes (see methods). The spliceosome complexes were then immunoprecipitated using the anti-SNRPB antibody (Y-12) followed by mass spectrometry analysis (Fig. 1C). The detailed list of identified peptides and proteins is shown in Table S1. The quantitative analysis of spliceosome-related proteins is shown as the heatmap in Fig. 1D. Several host splicing proteins showed distinct levels and nucleo-cytoplasmic distribution in the *Mtb*-infected macrophages compared to the uninfected cells (Fig. 1D). Thus, proteins like RPL4, SF3B2 and SYNCRIP showed contrasting nucleo-cytoplasmic distribution in the infected cells (Fig. 1D). In contrast, proteins like SNRNPs and HNRNPs showed severe depletion of the nuclear pool in the infected macrophages (Fig. 1D). The SNRNP proteins were particularly intriguing since they are critical for the assembly of small nuclear ribonuclear proteins (snRNPs)^23,26,40,41^. Dysregulation of these SFs suggests a systemic perturbation of the splicing machinery in *Mtb*-infected macrophages.

**Figure 1:**
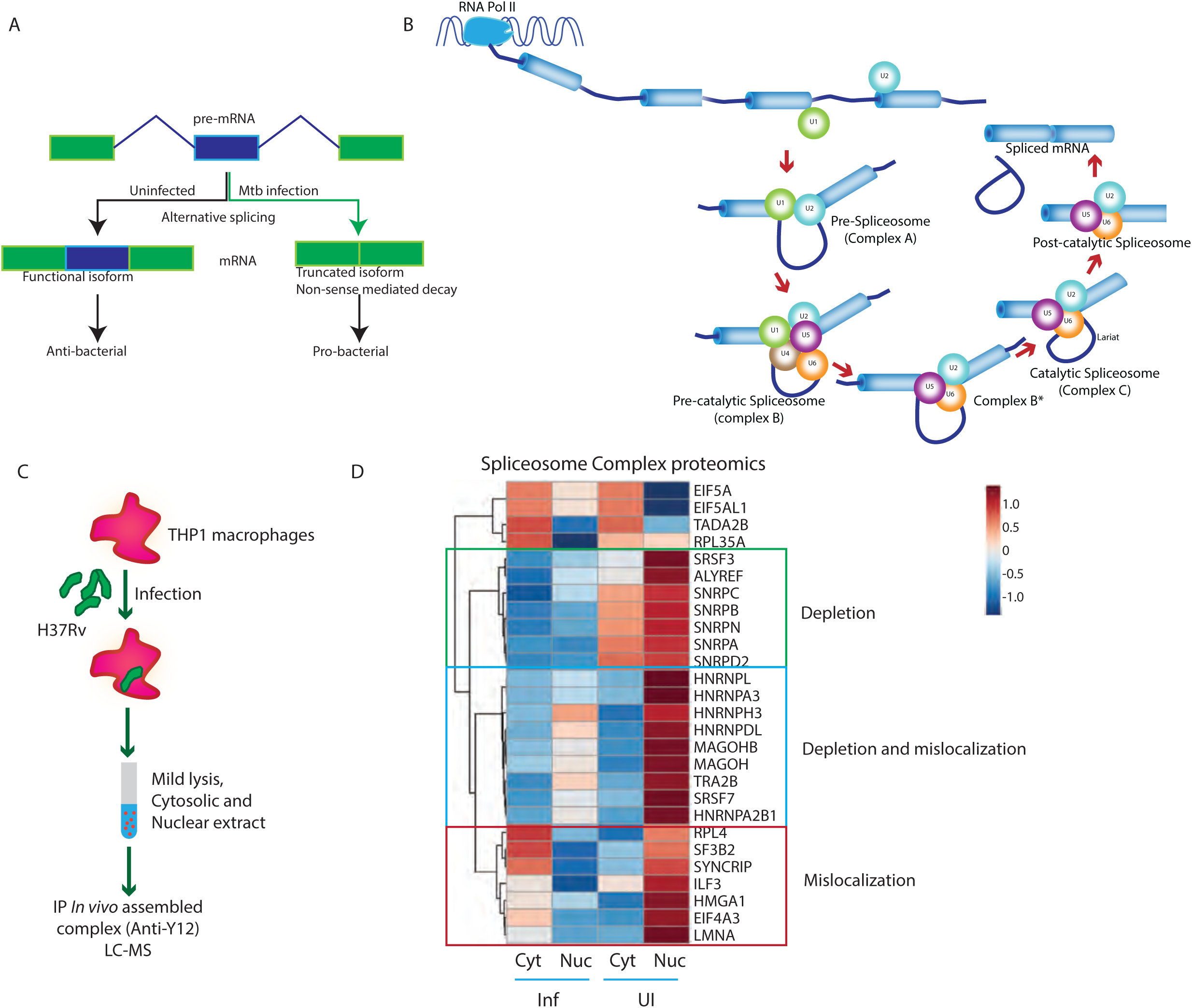
Dysregulation of host RNA splicing machinery in Mtb-infected macrophages. **A)** The schematic representation of perturbed alternative splicing resulting in exon skipping events upon *Mtb* infection. **B)** A schematic model depicting the ordered pathway of mRNA transcription by RNAP II and splicing catalysed by spliceosome complex. The U1snRNP binds the 5’ splice site and U2 snRNP binds the branch point adenine to form complex A. U4/U5.U6 snRNP binds to form complex B and causes conformational changes to form activated complex B. This is followed by the first transesterification reaction, releasing U1 and U4 snRNP and the second transesterification reaction forming complex C, releasing the lariat intron and U2, U5 and U6 snRNP. The U snRNP are the major players in splicing catalysis. **C)** The schematic of the spliceosome complex pulldown from the *Mtb* infected THP-1 macrophages 48 hours post-infection for Mass Spectrometry. **D)** Heatmap shows massive dysregulation in the level and cellular distribution of human proteins involved in RNA splicing and identified in MS of the spliceosome complex pulldown from the *Mtb* infected THP-1 macrophages 48 hours post-infection. The three boxes depict proteins that show depletion (green), depletion and mislocalisation (blue) and mislocalisation (red) patterns in Mtb-infected cells compared to the uninfected cells.

### Identification of *Mtb* proteins that could interact with host spliceosome complex proteins

Since a large number of spliceosome-associated proteins were dysregulated, we hypothesized that the virulent factors secreted by *Mtb* in the infected macrophages could likely be interacting with some of these proteins and responsible for their dysregulation. To test this hypothesis, we decided to screen whether any secreted protein from *Mtb* could interact with the components of host RNA splicing machinery using the yeast-2-hybrid (Y2H) method. The *Mtb* genes encoding the secreted proteins were part of the bait library (binding domain or BD vector), while the human spliceosome-associated proteins encoding genes constituted the prey library (activation domain or AD vector). In total, we had 384 *Mtb* genes in the BD library, including 37 negative controls (Table S2). For the secreted protein list, we relied mainly on our recent secretome study^42^ in addition to other *Mtb* secretome studies^36,43–45^ (Table S2). The AD library was comprised of the protein-coding isoforms of genes belonging to functional classes “RNA splicing” and “Spliceosome function” (∼162 clones), making it a unique custom library of human transcripts (Table S2). In the BD library, we also included a few negative controls (proteins which are never reported in the secretome analysis, Table S3). Four independent mating reactions between 384 BD clones and 162 AD clones, amounting to about 2,48,832, were conducted to obtain the interaction. The schematic for the Y2H assay is shown in Fig. 2A. In total, 123 Mtb proteins showed interactions with one or more SFs, together consisting of 439 incidences of interactions (Table S4). Out of these 439, 121 interactions were a result of autoactivation (Table S4). The BD clones showing autoactivation were excluded from the list of interacting proteins. A few negative control list clones also showed interactions, including 56 autoactivation incidences (Table S3). The interacting BD clones that did not show autoactivation were considered true interactors and mapped as a network to represent the *Mtb*-host spliceosome interactome, comprising 54 human SFs and 117 *Mtb* proteins (Fig. 2B, Table S5).

**Figure 2:**
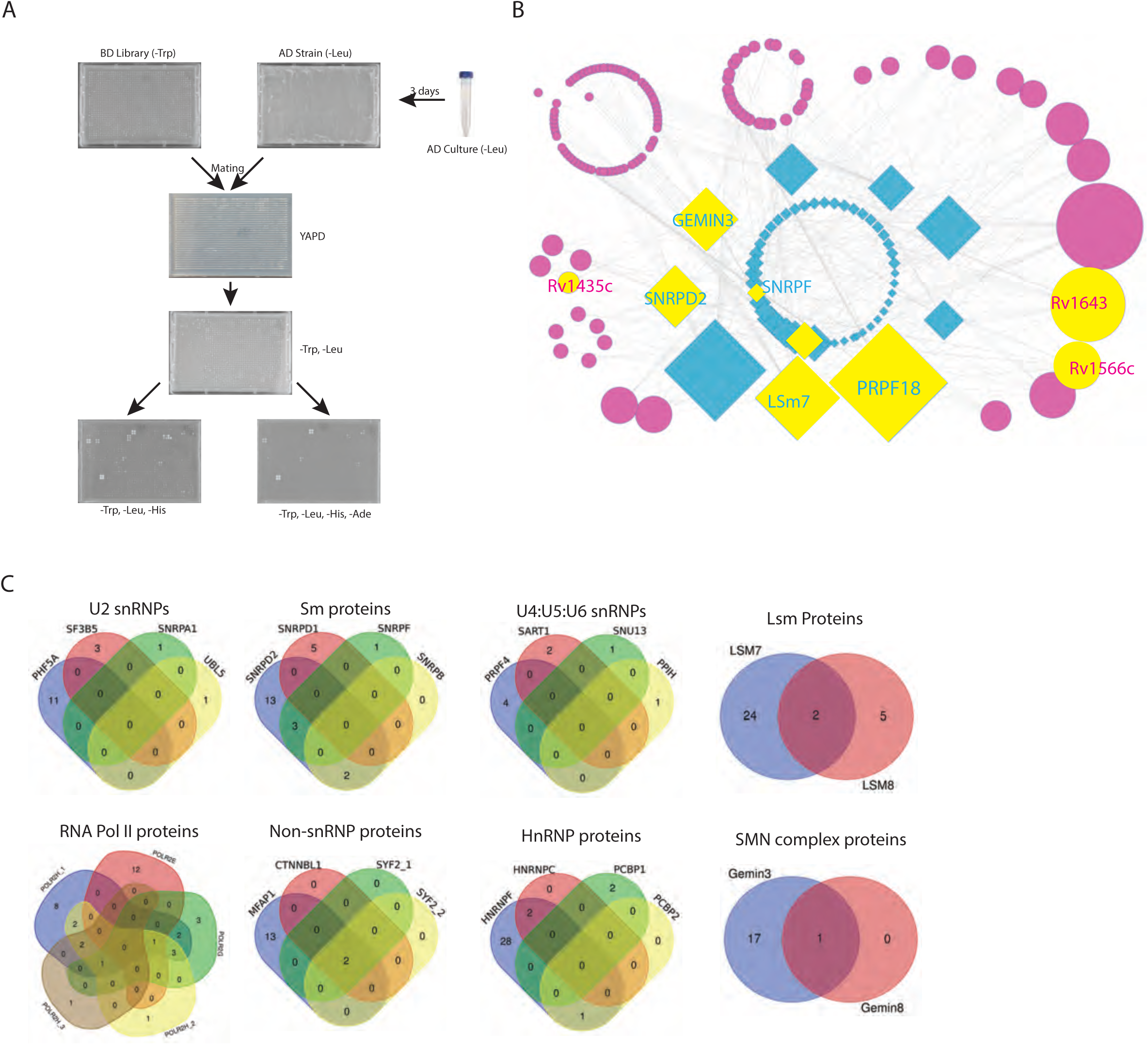
Mycobacterial secretory proteins and host spliceosome proteins interactome. **A)** The schematic flow chart for high throughput one-to-one Y2H assay in 1536 well format. Protein pairs showing interaction in 3 or 4 biological replicates were considered as positive interactions. B) *Mtb* proteins -host splicing factor interactome-the positive interactions from the Y2H screen were used to generate the network shown here. The diamond and circle represent the host and *Mtb* proteins, respectively, and the size corresponds to the number of interactions observed. The proteins coloured yellow in the network are described further in the subsequent sections. C) The Venn diagrams show proteins belonging to the major spliceosome components and their corresponding overlap with the interacting *Mtb* proteins.

The interactions spanned across all classes of host proteins that regulate RNA splicing, including RNA polymerases, Sm proteins, SMN complex proteins, U2 snRNP proteins, LSm proteins, U4/U5.U6 tricomplex proteins and non-snRNP proteins (Fig. 2C). The Venn diagram in Fig. 2C shows number of *Mtb* proteins interacting with proteins from each of these classes discussed above. Overall, host proteins from RNA Pol II subunits, Sm core, catalytic step II factors, HnRNPs and SMN complex and *Mtb* proteins from information pathway, virulence factors and conserved hypothetical proteins functional classes showed maximum interactions (Fig. S1A). Interestingly, host proteins from similar functional steps in RNA splicing shared their interacting partners from the *Mtb* (Fig. 2D). The Mtb proteins showing interaction also represented diverse functional classes (Fig. S1B). Finally, due to the inclusion of protein-coding splice variants of host SFs in the AD library, eighteen genes were represented by more than one transcript in the interactions list (Fig. S2). Out of those, POLR2H, SYF2, SNRPD2, LSm7, RBMX, LUC7L2, DCAF4, PPIH and U2AF1 showed interactions with the *Mtb* proteins (Fig. S2). Curiously, the spliced variants with a truncated coding region interacted with many *Mtb* proteins compared to their full-length counterparts (Fig. S2). While this observation suggests looking at protein homeostasis in the *Mtb*-infected macrophages more carefully, we decided to remain focused on our objective of understanding the mechanism of infection-induced alternative splicing.

### *Mtb* secretory protein Rv1435c could alter host RNA splicing

Several *Mtb* secretory proteins showed promising interactions in the Y2H assays above. Before taking any of these proteins for further mechanistic explorations, it was important to verify them on two accounts-one, if the identified *Mtb* secreted protein could alter host RNA splicing and two, whether the identified interacting partners in the Y2H assay also interacted with each other in a mammalian system. We picked up a few *Mtb* proteins, one showing a very high number of interactors (Rv1643, 22 interactions), another with an intermediate number of interactors (Rv1566c, 13 interactors) and a third with a low number of interactors (Rv1435c, 5 interactors) (Table S3). To test whether these *Mtb* proteins could alter host RNA splicing in a targeted manner, we first developed a few assays. For the functional assays, it was critical to choose target genes for which splicing events were to be measured. We chose around five target genes from the host, including *ACSL1*, *ACADVL*, *DDX11*, *DDX42* and *NFYA*, which consistently showed exon exclusion events upon *Mtb* infection^13^. We designed primers to score for the test exon exclusion event, exon inclusion event, and a set of primers from the exon junctions (Fig. S3A). The exon exclusion primers clearly showed, upon *Mtb* infection, an increase in the expression of selected transcripts of the genes where the test exon was excluded (Fig. S3B). On the contrary, with the exon inclusion primers, except for the *ACSL1* transcript, others either did not show any regulation (ACADVL and DDX42) or showed a decrease in expression upon *Mtb* infection (DDX11 and NFYA) (Fig. S3C).

Finally, primers designed from the junction of exons 1 and 3 conclusively showed that the selected transcripts showed clear cases of *Mtb* infection-induced exon exclusion (Fig. S3D). Next, we used a splicing reporter assay, which included a constitutive reporter construct *pRint* and a test construct *pGint* ^46^ (Fig. 3A). The *pGint* reporter construct was modified to include the test exon (middle exon) along with the flanking 100bp intronic sequences (Fig. 3A). The HEK293T cells stably expressing *pRint* were transfected with *pGint_ACADVL, pGint_ACSL1, pGint_DDX11, pGint_DDX42* and *pGint_NFYA* along with either a control non-interacting *Mtb* protein encoding plasmid (NIC) or plasmid encoding for *Rv1435c, Rv1566c or Rv1643* (Fig. 3B). Expressing NIC alone led to a marginal but consistent increase in the %pGint^+^ cells in the case of *DDX11* and *ACSL1* (Fig. 3C). Rv1435c and Rv1566c showed a significant increase in %pGint^+^ cells in the case of all except *ACSL1* over and above corresponding NIC expression (Fig. 3C). Unlike Rv1566c and Rv1435c, for Rv1643, significant increase compared to the NIC were noted in the case *ACADVL*, *DDX11* and *DDX42* but not in the case of *NFYA* (Fig. 3C). Despite the limitation that we used only a handful of target RNA splicing, we could show that the three selected *Mtb* proteins were able to alter host RNA splicing.

**Figure 3:**
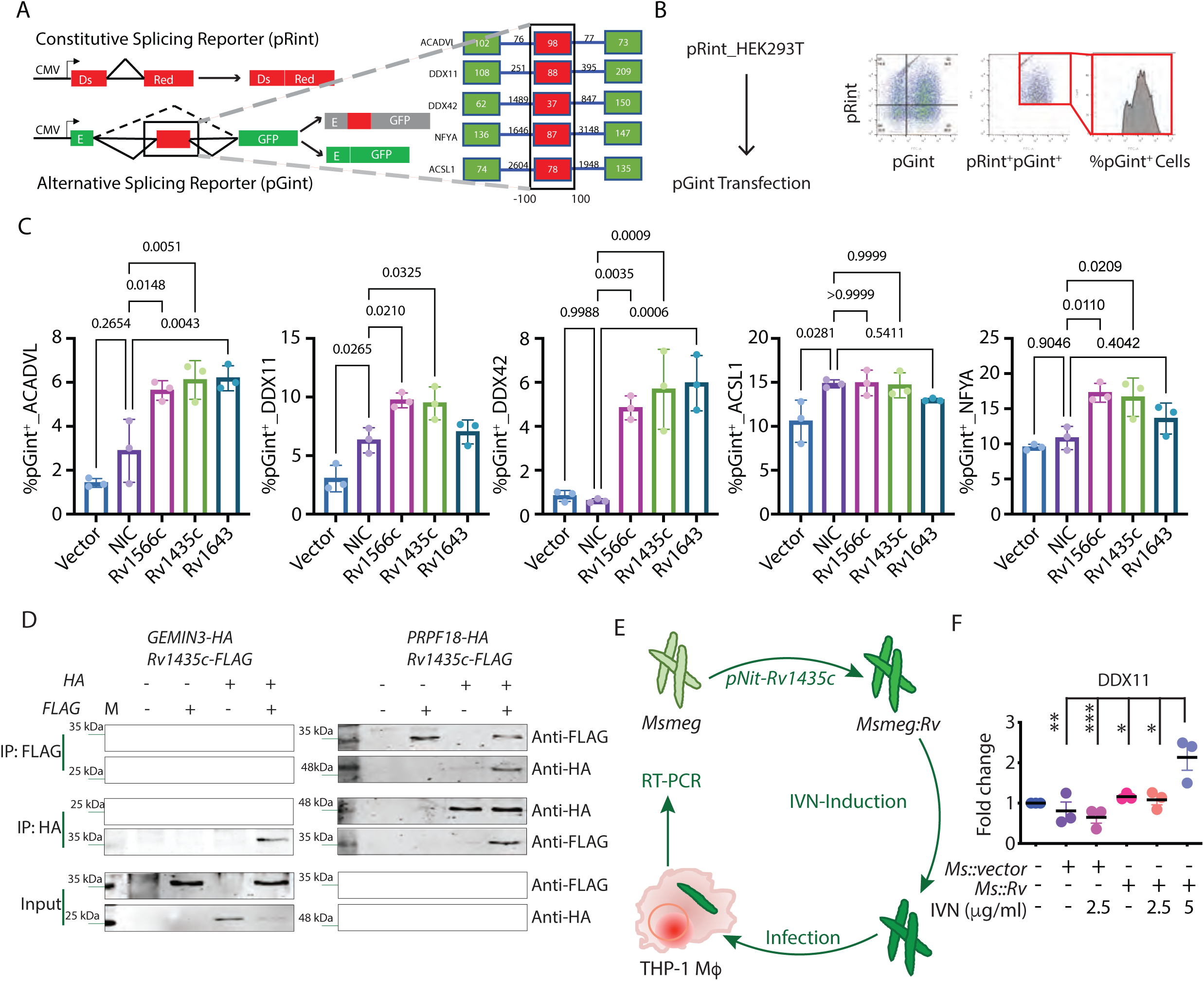
*Ex vivo* infection assays and reporter assays to test the role of *Mtb* proteins in host RNA splicing. A) Designing of the custom *in vivo* fluorescent splicing reporter. Schematic for the constitutive splicing reporter pRint with a short adenovirus intron intervening the DsRed gene with a strong 5’ and 3’ splice sites. In the pGint splicing constructs, the alternative exon of ACADVL, DDX11, DDX42, ACSL1 and NFYA, along with the flanking 100bp intronic sequences, were inserted within eGFP reading frame. The exon disrupts the reading frame, however, skipping of the test exon reconstitutes EGFP fluorescence. B) In cell reporter assay: the pRint expressing stable HEK293T cell lines were co-transfected with a pGint construct and pDest-FLAG construct encoding proteins Rv1252c (NIC), Rv1566c, Rv1435c, Rv1643, or empty vector control, followed by Flow cytometry. In the DsRed^+^ gate, the %FITC^+^ population was calculated to measure the extent of the test exon-skipping event. C) %FITC^+^ population for each of the indicated reporter constructs in the presence of indicated mycobacterial protein, non-interacting control, or vector controls. Each dot represents an independent biological replicate (N=3), indicating p values from One-way ANOVA. D) Validation of physical interactions between indicated pairs of Rv1435c proteins (FLAG-tagged) and host proteins – (HA-tagged) in HEK293T cells. Data is representative of two or more biologically independent transfection and pulldown experiments. E) The schematic representation of infecting THP-1 macrophages with *Msmeg* strains that express Rv1435c under IVN inducible promotor followed by RTPCT for assessing target gene splicing event. F) Isoform-specific qPCR to assess the impact of Rv1435c expressing *Msmeg* infection in THP-1 macrophages on exon skipping event in DDX11, N=3, indicating p values from One-way ANOVA.

In the above assay, among the three *Mtb* proteins analyzed, *Rv1566c* and *Rv1435c* showed a stronger influence on exon exclusion events. Among the two, Rv1435c also contains a signalP, which is essential for secretion through the classical bacterial secretory pathway ^43,44^. Secondly, while some function for Rv1566c was assigned earlier ^45,47^, Rv1435c remains an uncharacterised protein. Therefore, we decided to pursue further characterization of Rv1435c in regulating host RNA splicing events.

First, we verified the physical interactions between Rv1435c and its Y2H counterparts from the human SFs. Out of five interacting SFs, we picked up two Rv1435c interacting SFs, GEMIN3 or DDX20 and PRPF18, to verify interactions in the mammalian cells. Flag-tagged *Rv1435c* was co-expressed along with HA-tagged human counterparts in HEK293T cells, followed by co-immunoprecipitation. Consistent with the results from the Y2H screen, both anti-Flag and anti-HA pulldowns were able to co-immunoprecipitate each other, suggesting strong interaction of Rv1435c protein with the two human proteins GEMIN3 and PRPF18 (Fig. 3D). Therefore, Rv1435c was able to induce exon exclusion event in the *in-cell* reporter assay and also showed physical interaction with human SFs in the mammalian cells.

### A surrogate infection model using *M smegmatis* knock-in line shows a direct role of Rv1435c in altering host RNA splicing during infection

Next, to test whether Rv1435c could also impact these exon exclusion events during infection, we used the non-pathogenic *Mycobacterium smegmatis* (*Msmeg*) strain. We cloned Rv1435c under a Tet-inducible promoter and expressed it in the *Msmeg*, making it *Msmeg::Rv1435c* (Fig. 3D). We confirmed that Rv1435c was expressed and secreted by the modified *Msmeg* upon induction (Fig. S4A). Next, we infected THP-1 macrophages with *Msmeg, Msmeg::vector or Msmeg::Rv1435c,* which were either untreated or treated with ATC before infecting the cells (Fig. 3D). Specific exon exclusion events were then tested using RT-PCR assays against all the five target events described above. For *ACADVL*, *DDX42*, *ACSL1* and *NFYA*, while we did observe a trend, the increase in exon exclusion event when *Msmeg::Rv1435c* line was induced by anhydrotetracycline (ATC), it was not very consistent and significant (Fig. S4B). In the case of DDX11, infection with *Msmeg* alone, *Msmeg::vector* or *Msmeg::Rv1435c* did not induce the exon exclusion event (Fig. 3E). However, when *Msmeg::Rv1435c* was induced overnight with ATC, there was a significant increase in the case of *DDX11* exon exclusion event (Fig. 3E). Overall, results from the above sections show that Rv1435c from *Mtb* interacts with select host SFs and also alters specific host RNA splicing events.

### *Mtb* Rv1435c is a secretory protein with no assigned function

Once the role of Rv1435c in altering host RNA splicing was verified, we were curious to know what was known about this protein or its homologues from other species. The *Mtb* Rv1435c is a 202 amino acid-long polypeptide; a Pfam search revealed the presence of signalsP (1-33) at the N-terminal, indicating its secretion through the classic sec-mediated pathway (Fig. S4C). The remaining polypeptide segment (34-171) is classified as an extracytoplasmic region with a disordered region towards the C-terminal (172-202, Fig. S4C). Similar structural features were also revealed by AlphaFold Protein Structure Database^48^ (Fig. S4D). Sequence search revealed the presence of its orthologues in several pathogenic and non-pathogenic species of the genus *Mycobacterium* (Fig. S4E). Classifying these orthologues through a dendrogram analysis revealed an interesting pattern. In truly pathogenic species like *Mtb* and *M. marinum*, this gene is classified as a secreted protein with no function assigned (Fig. S4F). In opportunistic pathogenic species, like *M. helveticum, M. paratuberculosis, M. shinjukuense* and a few other unclassified species, this gene has been assigned the function of secreted beta-xylosidase (Fig. S4F). Even in a few truly pathogenic species like *M. lacus* and *M. heildelbergense*, this gene is characterized as secreted beta-xylosidase. It is possible that altering RNA splicing through interaction with the host spliceosome complex emerged as a function during the evolution of the pathogenic species.

### *MtbΔRv1435c* fails to alter target gene alternative splicing during infection

Next, to establish that Rv1435c could be directly involved in altering the host RNA splicing during *Mtb* infection, we deleted the *Rv1435c* gene in *Mtb* using recombineering^49^ to generate the knockout strain *MtbΔRv1435c* (Fig. 4A). The knockout (KO) of *Rv1435c* was verified by PCR using specific primers as described in the methods and also by RT-PCR to test the expression of Rv1435c RNA in the KO cells (Fig. 4B and 4C). We also generated an *Rv1435c* complemented strain by electroporating the WT *Rv1435c* encoding plasmid in the *MtbΔRv1435c* cells to generate *MtbΔRv1435c::WT*. Next, we infected THP-1 macrophages with the WT *Mtb*, *MtbΔRv1435c or MtbΔRv1435c::WT* and assayed for the exon exclusion events in the five target genes, as described earlier. While the WT *Mtb* infection, expectedly, increased the exon exclusion event for each of the five target genes, in *MtbΔRv1435c-* infected THP-1 macrophages, the exon exclusion events were significantly downregulated (Fig. 4C). In the THP-1 macrophages infected with *MtbΔRv1435c::WT*, the exon exclusion event was restored to the levels observed during WT *Mtb* infection (Fig. 4C). These results indicate that Rv1435c protein of *Mtb* can alter the targeted host RNA splicing events in the infected macrophages and therefore, we named it as **h**ost **s**plicing **r**egulator 1 or hsr1.

**Figure 4:**
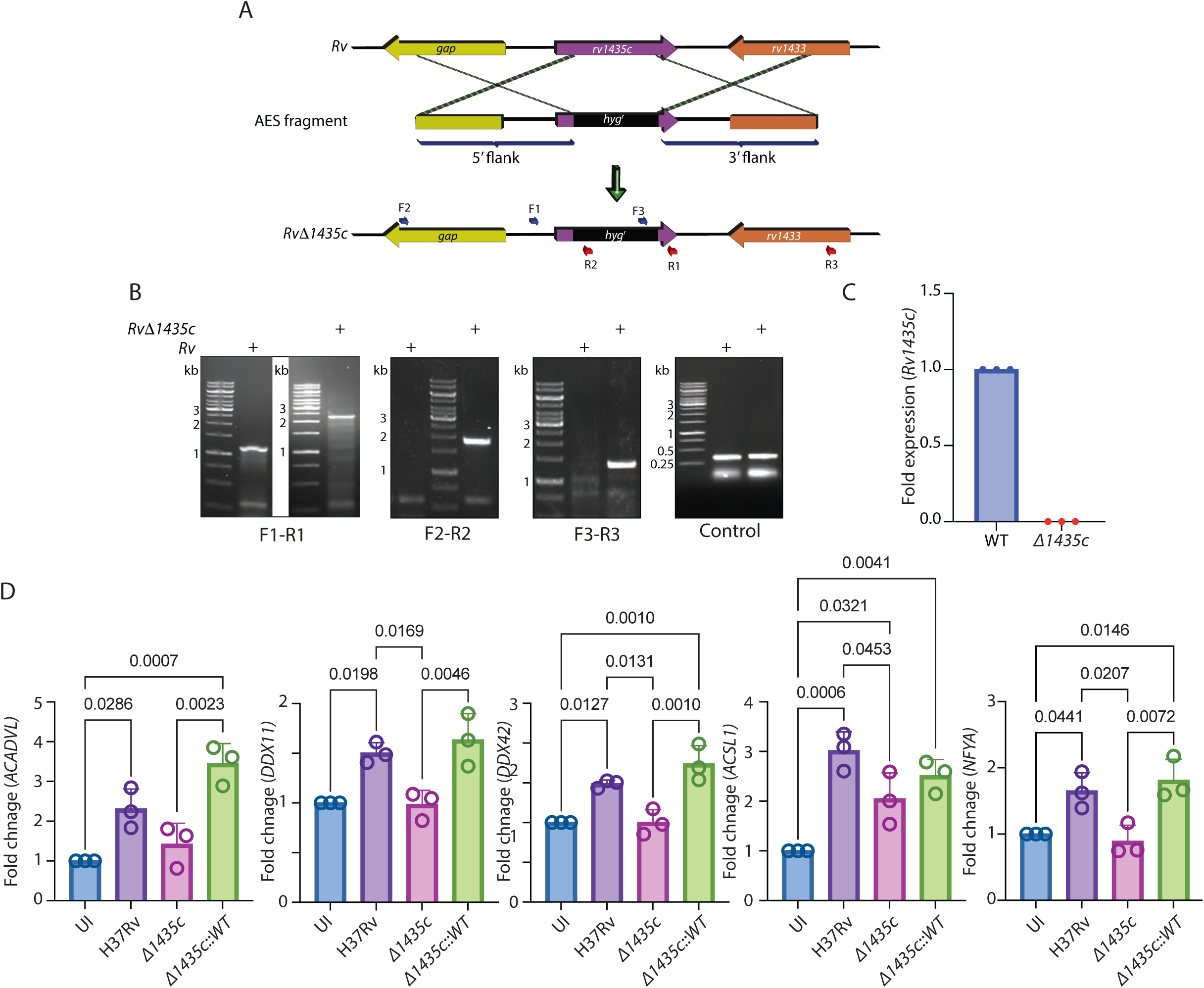
Role of Mtb Rv1435c protein in altering host RNA splicing. A) The schematic of knocking out the *Rv1435c* or *hsr1* gene in H37Rv using the recombineering approach. Positive colonies were screened with the primers F1-F3 (forward) & R1-R3 (reverse), as shown by the arrows. B) Agarose gel electrophoresis to confirm the replacement of 609bp product (*Rv1435c*) with 1.4Kb hygromycin cassette in the knockout line (F1-R1), absence of hyg in the WT (F2-R2 and F3-R3) and control segment (*rbpA*). C) Q-PCR to show the loss of expression of *Rv1435c* in the knockout strain D) Q-PCR for the target transcripts as mentioned after infection with H37Rv, H37Rv*Δhsr1* and H37Rv*Δhsr1::WT* strain. Data from three biological replicates (N=3), indicating p-values from a one-way ANOVA analysis.

### *Mtb* hsr1 binds to and sequesters U5snRNA in the cytosol

Next, we wanted to understand the mechanism of hsr1-mediated alteration in host RNA splicing. In the spliceosome complex pulldown results (Fig. 1) and the Y2H screening results (Fig. 2), several proteins from the snRNP complexes were present. This included proteins from the SMN complex like GEMINs, DDX proteins and those from the heptameric core that are recruited to the 3’ sm Site of snRNAs for the biogenesis of snRNPs. The biogenesis of snRNPs follows an evolutionarily conserved nucleo-cytoplasmic shuttling, where snRNAs, upon being synthesized in the nucleus, are exported to the cytosol (Fig. 5A). In the cytosol, snRNAs bind with the SMN complex, followed by the recruitment of sm Core proteins^50^ (Fig. 5A). We anticipated that snRNP biogenesis, due to its cytosolic trajectory, could be a potential target of *Mtb* proteins-mediated alteration. We analyzed the levels of two snRNAs-U2 and U5 snRNAs, in the *Mtb*-infected macrophages using Fluorescence In Situ Hybridization (FISH). While the U2 snRNA showed no noticeable pattern (Fig. S5A), we noted a significant regulation in U5 snRNA levels in *Mtb*-infected macrophages at 24 and 48 hpi (Fig. 5B). While there was a significant decline in U5 snRNA levels at 24 hpi, we noted a significant increase in U5 snRNA levels at 48 hpi (Fig. 5B-C). To our surprise, we also observed a direct interaction between *Mtb* phagosomes and U5 snRNA (Fig. 5D). The U5 snRNA interaction with the *Mtb* phagosomes increased over time from almost no noticeable interaction at 24 hpi to significantly high interaction at 48 hpi (Fig. 5D). It appeared that *Mtb* phagosomes were binding and sequestering the U5 snRNA in the cytosol (Fig. 5D). Next, to test whether hsr1 was by anyway involved in this binding, we infected THP-1 macrophages with WT *Mtb* or *Δhsr1* and quantified the FISH signals to compare the interaction between U5 snRNA and phagosomes (Fig. 5E). We noted a significant decline in U5 snRNA interaction with the phagosomes in the case of *Δhsr1* infection (Fig. 5E). The above results suggest that hsr1 was responsible for binding with U5 snRNA in the cytosol. Any defect in snRNP biogenesis is known to alter the global RNA splicing pattern^16,30,31,51–54^. Therefore, it was likely that the altered RNA splicing events regulated by hsr1 could be due to its interaction with the U5 snRNA.

**Figure 5:**
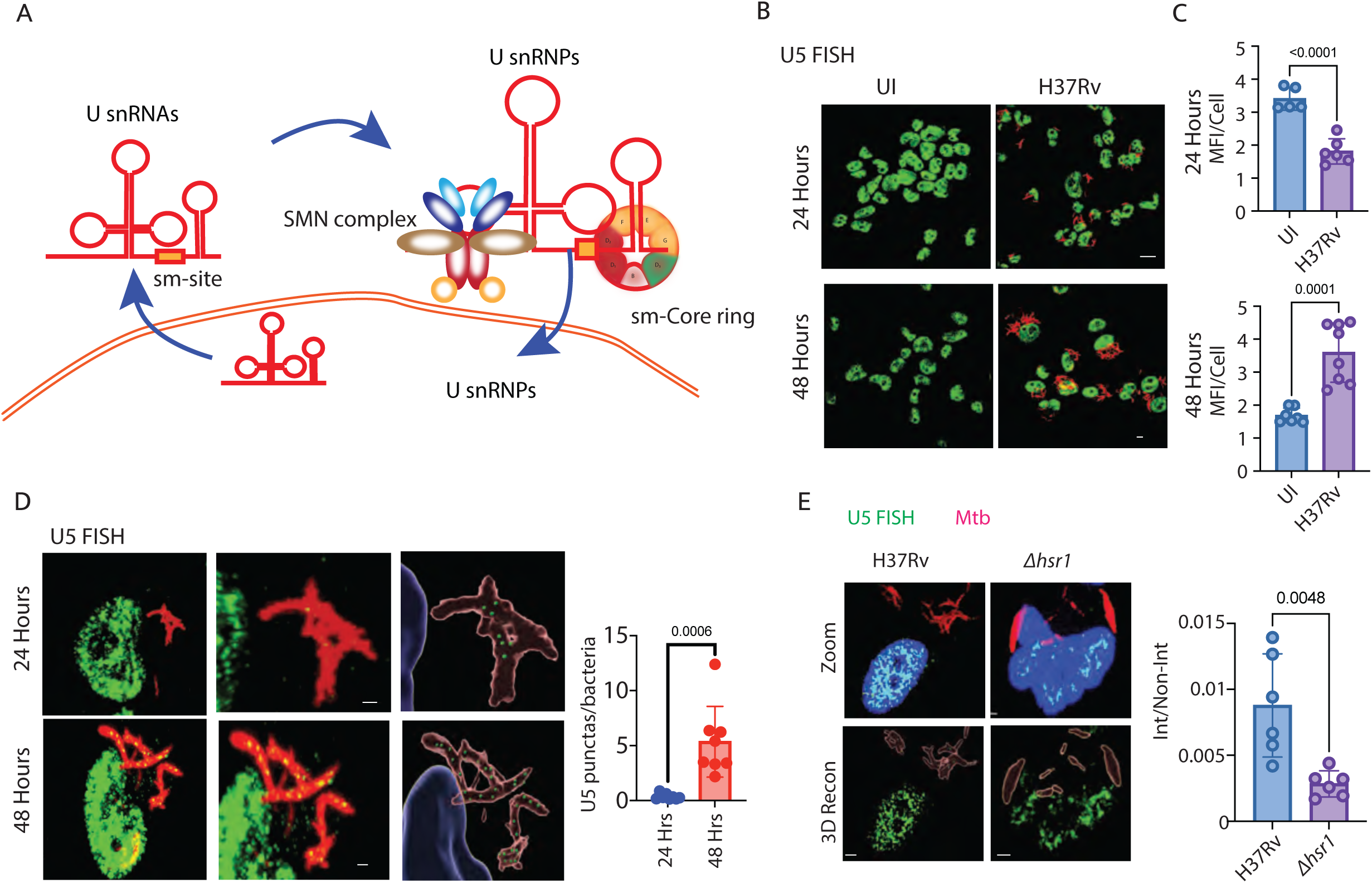
Direct interaction of U5 snRNA with Mtb phagosomes. A) Schematic representation of U snRNA nucleo-cytoplasmic trajectory and assembly of Sm core ring on snRNA by SMN complex to form snRNPs B) Representative images for RNA FISH against U5 snRNA in THP-1 macrophages upon *Mtb* (mCherry-H37Rv) infection at 24- and 48-hrs post-infection. The U5 snRNA probe was labelled with Alexa 488 (green), and the nucleus was stained with DAPI (blue). C) Plots show quantification of U5 snRNA FISH MFI in uninfected and *Mtb*-infected macrophages at 24 ad 48 hpi. D) Direct interaction between H37Rv (red) and U5snRNA at 24- and 48-hrs post-infection. U5snRNA puncta within 0.2 microns from the H37Rv surface are shown as green dots on the 3-D constructed Mtb contours. The 3D reconstruction of the nucleus, U5 snRNA foci and bacteria (see methods) was done to show the U5 snRNA Foci at <0.2 microns distance from the bacterial phagosomes. The quantification of interacting U5 snRNA punctas is shown in the plot at the right. Scale bar: 2 μm. E) Representative image for the U5snRNA FISH in THP-1 macrophages infected with mCherry expressing WT H37Rv or H37Rv*Δhsr1* strains. The plots at the right show the ratio of interacting to non-interacting U5snRNA with WT H37Rv or H37Rv*Δhsr1*. Data is from three biological experiments, two fields each, representing p-value from a Student’s t-test analysis.

### Mtb *hsr1* interacts with SNRPF, a key sm Core protein in the cytosol

The hsr1-dependent interaction of *Mtb* phagosomes with U5 snRNA raised the question of whether the interaction involved only the U5 snRNAs or reflected the binding of the entire U5snRNP complex. The snRNP complexes, as discussed above, comprise proteins from SMN complexes like GEMINs, SMNs, etc and the heptameric ring of sm Core proteins, including SNRPB, SNRPD1, SNRPD2, SNRPC, SNRPF, etc. We analyzed some of the marker proteins of snRNPs using confocal microscopy, including DDX23, GEMIN8, GEMIN3, Lsm7, SNRPD2 and SNRPF (Fig. 6A and S6). While some of these, like DDX23, GEMIN8, SNRPD2 and Lsm7, showed some random interactions with the *Mtb* phagosomes, none of them showed a pattern similar to that of U5 snRNA (Fig. S6). The only protein which showed U5 snRNA-like interaction with *Mtb* phagosome was SNRPF (Fig. 6A). Moreover, we further verified that SNRPF interaction with *Mtb* phagosomes was also hsr1 dependent since there was a sharp decline of their interaction when *Δhsr1* strain was used for infection (Fig. 6B). Curiously, SNRPF was also part of our Y2H prey library; however, we did not see any interaction between SNRPF and hsr1 in the Y2H assay. Since our criteria for selecting positive interactions were very stringent, we sought to test whether these two proteins interact in the coimmunoprecipitation assay. In the HEK293T cells overexpressing Rv1435c-FLAG and SNRPF-HA, we observed that FLAG immunoprecipitation was able to pulldown SNRPF-HA (Fig. 6C). However, since other smCore proteins line SNRPD2 did not show noticeable interaction with *Mtb* phagosomes, we believe that Mtb phagosomes, though hsr1 specifically binds with U5 snRNA and SNRPF. These results suggest that *Mtb* hsr1 facilitates the phagosome interaction with U5 snRNA and the smCore protein SNRPF, likely contributing to an altered host RNA splicing.

**Figure 6:**
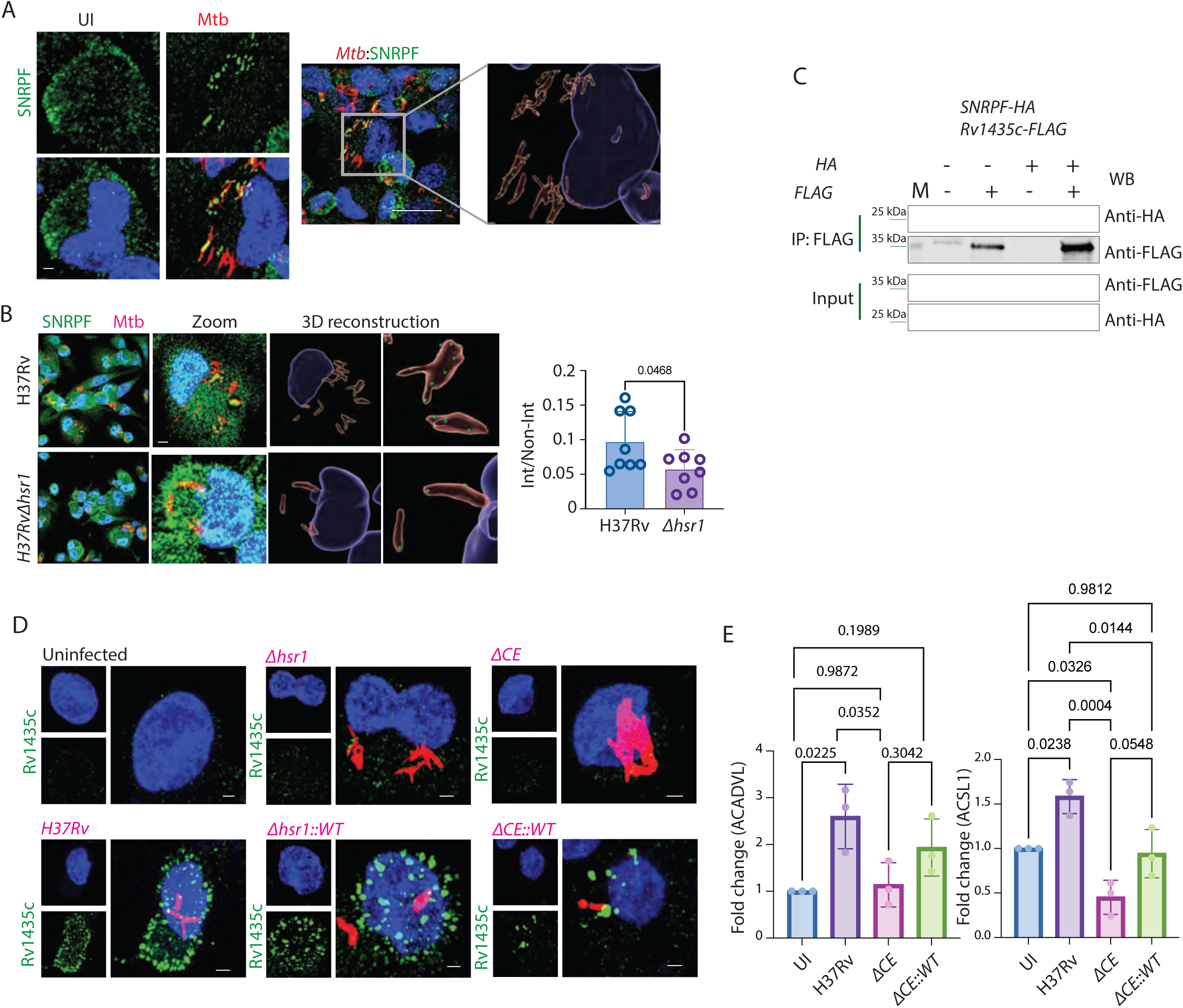
*Mtb* hsr1 secretion is ESX1-dependent, and impacts host RNA splicing by sequestering SNRPF. A) Image showing SNRPF colocalising with bacterial phagosomes. The 3D reconstruction of the nucleus, SNRPF foci and bacteria (see methods) was done to show the SNRPF Foci at <0.2 microns distance from the bacterial phagosomes. Scale bar: 2 μm and p values indicate Student’s t-test. B) Representative images of SNRPF staining in THP-1 macrophages infected with WT H37Rv or H37Rv*Δhsr1* strains. For the plot below, the number of SNRPF puncta within 0.2 microns from the H37Rv surface was counted as interacting and shown as the ratio of interacting to non-interacting SNRPF with WT H37Rv or H37Rv*Δhsr1*. Data is from three biological replicates with two or more fields per slide, representing p-values from Student’s t-test. C) Validation of physical interactions between Rv1435c (FLAG-tagged) and SNRPF (HA-tagged) in HEK293T cells using co-immunoprecipitation. Data is representative of two or more biologically independent transfection and pulldown experiments. D) Representative images of THP1-macrophages stained with DAPI (nucleus) and Rv1435c antisera (green) upon infection with mCherry expressing WT H37Rv, *Δhsr1, Δhsr1::WT, ΔCE* and *ΔCE::WT* strains. Images are representative of two independent biological experiments. Scale bar-2 μm. E) The exon-skipping event in ACADVL and ACSL1 transcripts upon infection with WT H37Rv, H37Rv*ΔCE* and H37Rv*ΔCE::WT* at 48hpi from three biologically independent experiments. The p values are from One-way ANOVA.

### *Mtb* hsr1 secretion and, therefore, its impact on host RNA splicing is ESX1-dependent

The hsr1 protein was a part of the list of *Mtb* proteins present in the culture filtrate, indicating their secretion ^42^. However, whether hsr1 also gets secreted during macrophage infection is unknown. Secretion during infection could be a critical factor for the ability of hsr1 to interact with the human spliceosome components. We generated antisera against hsr1 and confirmed its specificity by immunoblotting and microscopy before using it to examine the secretion of hsr1 during macrophage infection (Fig. S6A). In the uninfected macrophages, expectedly, we could not see any signal for hsr1; however, in the WT *Mtb*-infected macrophages, we could see several hsr1 foci on *Mtb* phagosomes and in the cytosol (Fig. 6D). When we used the Δ*hsr1,* there was no noticeable signal of hsr1 protein in the phagosomes or in the cytosol (Fig. 6D). When we used the complemented strain Δ*hsr1::WT,* the signal for hsr1 on the bacterial phagosome and in the cytosol was restored (Fig. 6D). We next question, how *Mtb* hsr1 was secreted during macrophage infections. For access to cytosol, the type VII secretion system ESX is critical, which is known to form pores in the phagosome membrane, allowing access to the cytosol^37,45^. To test that, we used a strain lacking CFP10 and ESAT6 (*ΔCE*), the two critical components of the ESX1 secretion system ^42,55,56^. In *ΔCE-*infected macrophages, we could not observe any signal for hsr1 in the cytosol of infected cells (Fig. 6D). Moreover, even secretion of hsr1 appeared to be ESX1-dependent since there was no hsr1 signal from the phagosomes (Fig. 6D). This was surprising since sequence analysis predicted a putative signalP sequence at the N-terminal of hsr1, making it a substrate of sec system^42^. Finally, when we used a complemented strain *ΔCE::WT*, where both ESAT6 and CFP10 were restored, we could see hsr1 signal, both at the phagosomes and in the cytosol (Fig. 6D). These results suggest that ESX1 pathway plays a central role in hsr1 secretion. The logical question next was to test whether that also impacts exon exclusion events in the host target genes through alternative splicing. We compared specific exon exclusion events in the case of ACADVL and ACSL1, and in both cases, *ΔCE* strain infection behaved similar to *Δhsr1* infection, whereas *ΔCE::WT* complemented strain showed exon exclusion similar to the WT *Mtb* or *Δhsr1::hsr1* infection (Fig. 6E). With the above results, we conclude that hsr1-mediated impact on host RNA splicing was dependent on its secretion in ESX1-dependent manner.

### *Mtb* hsr1 is a virulence factor and acts by sequestering SNRPF

Results so far show that without hsr1, *Mtb* could not alter the splicing events in the manner caused by the WT strain. We wondered whether this loss of function also impacted the intra-macrophage trajectory of *Mtb*. In *ex vivo* infected THP-1 macrophages, the growth of the *Δhsr1* strain was markedly reduced compared to the WT *Mtb* (Fig. 7A). In agreement with that, *Δhsr1* strain infected macrophages showed increased Cathepsin D staining and also an increase in the co-localization of *Mtb* with Cathepsin D, compared to the WT *Mtb* (Fig. 7B). This suggests that *Δhsr1* strains fail to execute the phagosome maturation arrest as is known for the WT *Mtb* ^57^. Since hsr1 was also responsible for binding with SNRPF in the cytoplasm of THP-1 macrophages, we next overexpressed SNRPF in THP-1 macrophages and infected them with WT *Mtb*, expecting that stoichiometric abundance of SNRPF should overcome the impact of hsr1 during WT infection and would help better control of bacterial growth. In THP-1 macrophages expressing SNRPF and infected with WT *Mtb*, we noted increased MFI for SNRPF and a reduction in the bacterial CFU (Fig. 7C). No such effect was seen when another sm Core protein SNRPD2 was overexpressed in these cells (Fig. S6B). We next compared the in vivo infection of Wt and *Δhsr1* strains in the mice model (Fig. 7D). Upon infection in mice, the *Δhsr1* strain was significantly compromised in the establishment of infection compared to the WT *Mtb*, as reflected by a reduced bacterial burden in the lungs of infected mice at 2- and 4-weeks post-infection (Fig. 7E). We also analyzed SNRPF immunostaining in the lung sections of uninfected WT *Mtb*-infected and *Δhsr1-*infected mice with anti-SNRPF antibody. Consistent with the results observed during *ex vivo* infection in THP-1 macrophages, the lung sections from WT *Mtb*-infected mice showed significant accumulation of SNRPF in the extranuclear space, suggesting binding and possible sequestration in the cytoplasm (Fig. 7F). This cytosolic staining of SNRPF was not observed in the lung sections from mice that were infected with *Δhsr1* strain (Fig. 7F). Thus, cytosolic sequestration of SNRPF by *Mtb* protein hsr1 is an important virulence mechanism of *Mtb*.

**Figure 7:**
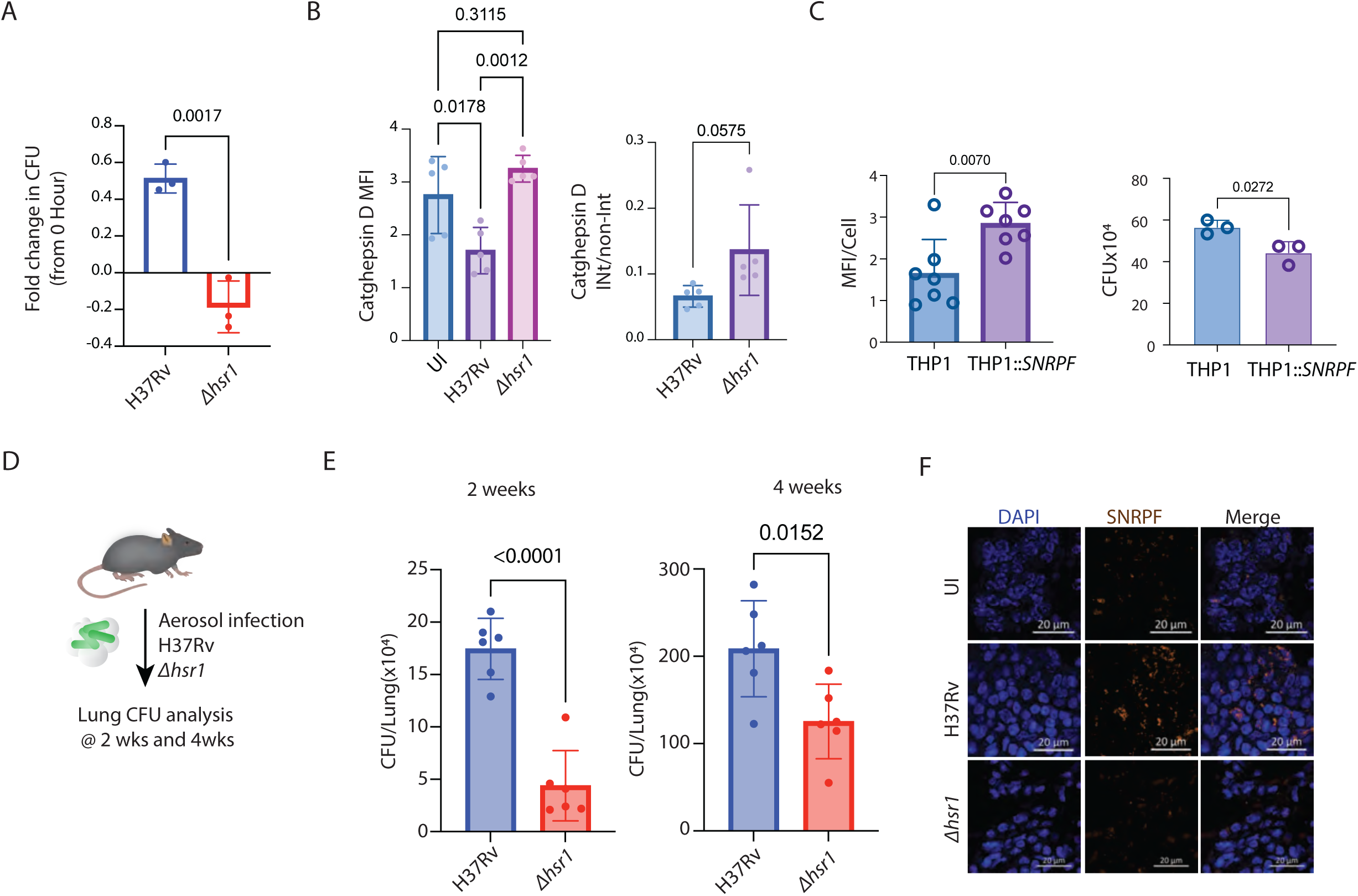
Regulation of host alternative splicing as a virulence mechanism. A) THP-1 macrophages were infected with WT H37Rv or *Δhsr1* strains. At 48 hours post-infection, bacterial load was examined using CFU plating. Data represent fold change in CFU at 48 hrs compared to 0 hr post-infection. (N=3), p values are from Student’s t-test. B) The plot shows quantification for staining of cathepsin D in THP1 macrophages infected with WT H37Rv and *Δhsr1*at 48hpi. The different parameters plotted show overall MFI (MFI/cell) and the ratio of cathepsin D foci interacting with bacterial phagosomes (int/non-int). Each dot represents a field analyzed, and an average of a hundred cells are quantified, representing p-values from one-way ANOVA (MFI) or Student’s t-test (Int/non-Int). C) The plot shows quantification (MFI/cell) for staining of SNRPF in WT THP1 macrophages and THP1*::SNRPF* infected with WT H37Rv at 48hpi. Each dot represents a field analyzed, and an average of a hundred cells are quantified. The plot on the right shows the bacterial survival after 48 hours post-infection in WT THP1 and THP1*::SNRPF*. Each dot represents an independent biological experiment. Indicated p values represent Student’s t-test. D) Schematic for study design using mice model of infection E) BALB/C mice (6-8 weeks old) were infected with 200 CFU of WT H37Rv and *Δhsr1* in separate experiments. At 2- and 4-weeks post-infection, bacterial load in the lungs of the animals was determined by CFU plating. N=6, representing p-values from Student’s t-test. F) The 4-week post-infection lung tissue sections of uninfected, WT H37Rv and Δhsr1 were stained with DAPI (nucleus) and anti-SNRPF antibody. Scale bar: 20 μm

### Altered RNA splicing in monocytes from human TB subjects and distinct SNRPF dysregulation in the human TB granulomas

To understand the dysregulation of RNA splicing in human TB subjects, we re-analyzed RNAseq data from the monocytes of human pulmonary TB subjects and corresponding healthy contacts^58^. The dataset comprised monocyte transcriptomes from 20 PTB subjects and 19 healthy controls. We analyzed this data set for transcript-level quantification (see methods, table S6). Using all the significantly regulated transcripts, we could observe distinct clustering of PTB subjects and healthy controls in the PCA analysis (Fig. S6C). Among the large number of transcripts showing significant regulation, we focused specifically on transcripts belonging to functional classes like RNA splicing, metabolism, immune response and inflammation. As can be seen from the heat maps in Fig. 8A, based on the level of transcript expression within each of the functional classes, the PTB group could be distinctively identified from the healthy controls. This remarkable observation highlights systemic RNA splicing alterations in tuberculosis patients. Finally, to assess whether an SNRPF-dependent mechanism plays a role in altering RNA splicing in human subjects, we screened human intestinal TB (ITB) granulomas for SNRPF by immunohistochemistry.

**Figure 8:**
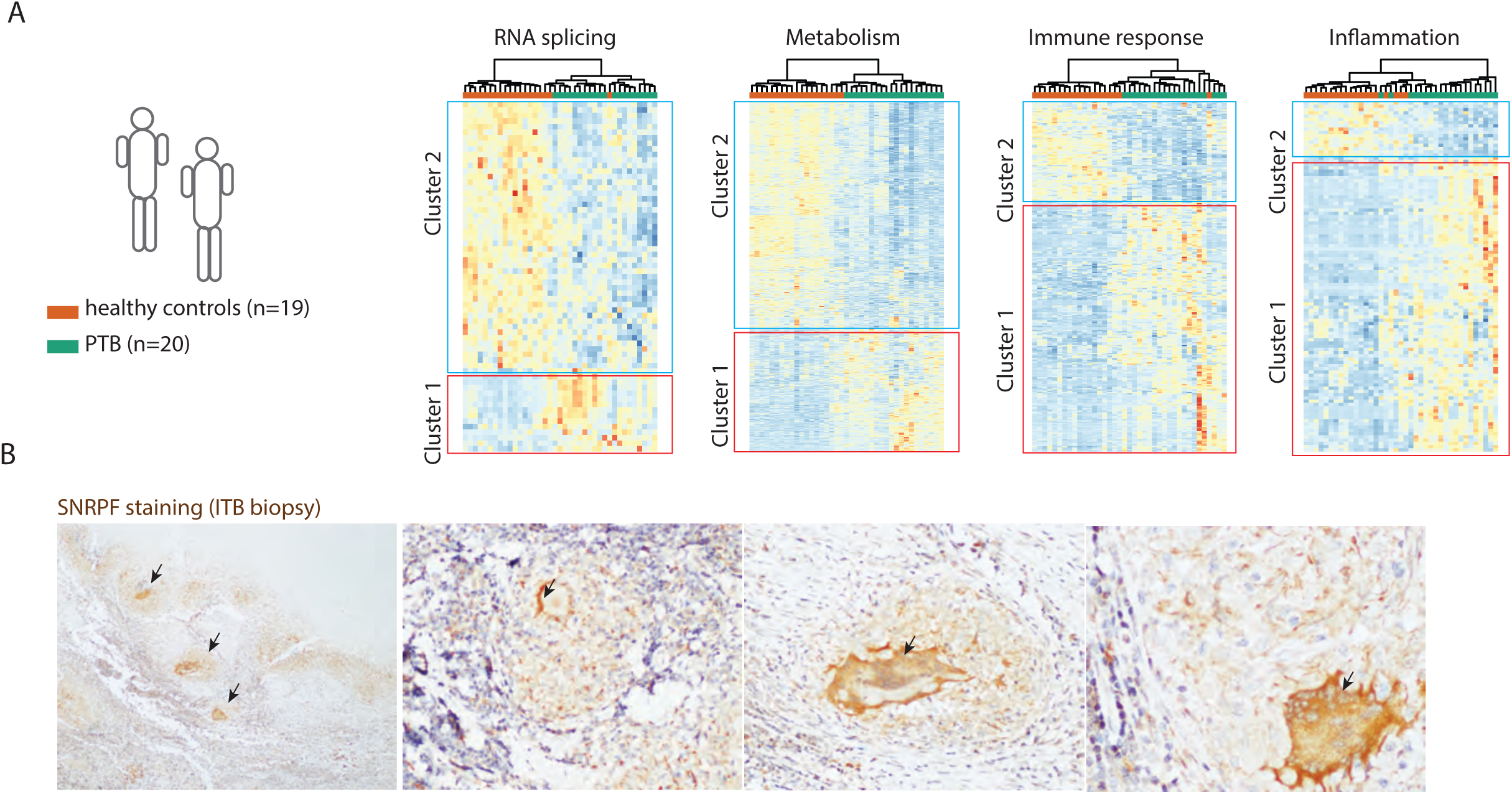
Altered RNA splicing and distinct SNRPF dysregulation in the granulomas of human TB subjects. A) RNAseq data from the monocytes of pulmonary TB cases (N=20) and healthy contacts (N=19) was analyzed to assess the expression of differential alternatively spliced transcripts. Genes belonging to the indicated functional classes were selected for the cluster analysis. Cluster 1 in each case shows transcripts upregulated in PTB cases. B) Biopsy sections from human intestinal tuberculosis granulomatous lesions were stained with anti-SNRPF antibody. The arrows indicate multinucleated giant cells with intense staining for cytosolic SNRPF. Black arrows show areas of intense SNRPF staining in the granulomas and in the giant cells.

**Figure 9:**
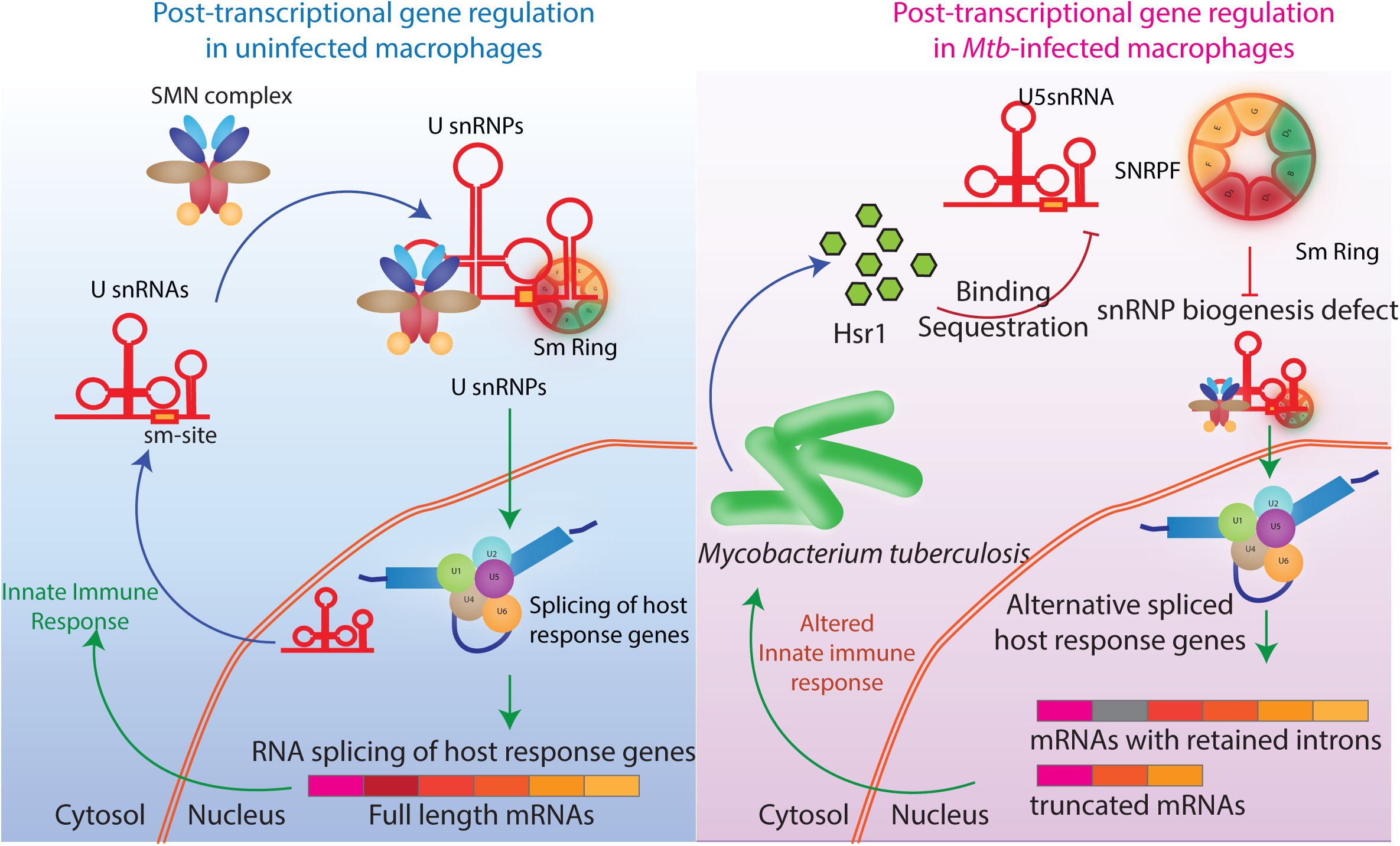
A model to show *Mtb*-mediated alteration of host RNA splicing by blocking the snRNP biogenesis and post-transcriptional gene regulation in the macrophages.

Consistent with the earlier results from *ex vivo* infection in THP-1 macrophages and *in vivo* infection in mice, the human ITB granulomas showed extensive SNRPF staining (Fig. 8B). Moreover, these staining were more pronounced in the multinucleate giant cells and neighbouring histiocytes (Fig. 8B and S6D). Additionally, we could observe strong cytosolic staining in the granulomas (Fig. 8B). No such distinct staining of SNRPF was noted in the control tissues (Fig. S6E). Similarly, there was no specific staining difference for another sm Core protein SNRPD2 between the control and ITB sections, again underscoring the specificity of the SNRPF-mediated mechanism (Fig. S8). Taken together, using results from *ex vivo* infection, *in vivo* infections and from the human TB subjects, we conclude that SNRPF sequestration and consequent alteration in the host RNA splicing aids bacterial growth and survival within the host, thereby contributing to its pathogenicity.

## Discussions

The ability of *Mtb* to adapt to the hostile immunological environment in the hosts is remarkable. Almost every physiological process relevant to host immune responses appears to be targeted by the *Mtb,* enabling the pathogen’s survival and consequent pathology. This study specifically looked at RNA splicing, a post-transcriptional gene regulation mechanism in the eukaryotes, to understand how *Mtb* achieves the targeted impairment of the seemingly complex process of RNA splicing in the host. As shown earlier by us and others, infection-induced altered RNA splicing helps the bacteria control host responses by preventing the synthesis of target genes in a fully functional form ^13,15,59–61^. Considering that RNA splicing is fundamental to eukaryotic gene expression, which in turn is critical for shaping the cellular responses to infection, the mechanistic understanding of this process could have a transformational impact on our ability to develop novel strategies towards controlling TB.

Since RNA splicing is a multi-step process involving the assembly of a large multi-molecular complex, our working hypothesis was that certain secreted proteins from *Mtb* could interact with and, therefore, interfere in the assembly of spliceosome complexes, thereby altering the pattern of RNA splicing. Proteomic analysis of the spliceosome complexes from the infected macrophages and the Y2H screen between bacterial secretome and host SFs, through independent approaches, did converge to some dysregulation at the level of snRNPs, the key components of spliceosome complexes. The assembly of snRNPs require snRNAs to get transported to the cytoplasm, where they interact with the components of the SMN complex and the heptameric ring called sm Core proteins, together forming the snRNPs, that enter the nucleus and participate in the RNA splicing reactions^23,24,31,50,53^. While the interaction network obtained by Y2H was dense and practically impossible to verify all those interactions through independent techniques, we rationally chose a few *Mtb* genes identified through the Y2H screen for further validation. With the customized assays developed for the purpose, we narrowed down on Rv1435c or hsr1, as the major regulator of host RNA splicing during infection. Apart from our observation of its ability to alter host RNA splicing, there is no defined function of Rv1435c in the literature, making it an attractive target for mechanistic explorations.

Interestingly, the orthologues of Rv1435c in other mycobacterial species are classified as secreted xylosidase, an activity lacking in the *Mtb* hsr1. Mechanistically, hsr1 allowed *Mtb* to bind in the cytosol with two key components of snRNPs, U5 snRNA and SNRPF.

Experiments with SMN complex proteins like GEMIN3 or GEMIN8, smCore proteins like SNRPD2 and snRNAs like U2 snRNA reveal that this interaction was most likely selective with U5 snRNA and SNRPF, and it did not represent the interaction of *Mtb* with whole snRNP complex. Through studies in unrelated systems like neurodegenerative diseases and cancer, it is shown that stoichiometric disbalance of snRNP components like snRNA, SMN complex proteins or smCore proteins can dramatically alter the snRNP complex biogenesis and impact RNA splicing^30,31,34,50,54,62–64^. U5 snRNA is classically known for its role in recognition of the 3’ splice site and, therefore, under depleting U5 snRNA condition, there could be a fierce competition between various 3’ acceptor splice sites, causing weaker splice junctions to lose out and not get incorporated into the final processed mRNA.

Similarly, SNRPF is part of several snRNP complexes, and its dysregulation due to cytoplasmic binding with *Mtb* may have a widespread impact on RNA splicing ^31,54,63^. Crucially, the nucleo-cytoplasmic shuttling of snRNA for snRNP biogenesis is an evolutionarily conserved quality control mechanism^53^. However, it appears that *Mtb*, through its course of evolution, learnt to disrupt this process without actually going to the nucleus.

The snRNP biogenesis defect appears to be operational in the lungs of *Mtb*-infected mice and the human TB granuloma sections. Since human PTB subjects also showed massive alterations in the RNA splicing pattern, to the extent that they could classify healthy from the PTB subjects, it is plausible that this mechanism could cause systemic RNA splicing defect, underlying many pathological states.

Interestingly, very limited reports highlight bacterial manipulation of host RNA splicing ^13,15,65^. A large number of studies focus on chronic pathology like neurodegenerative disorders or cancer^51,64^. Among infectious diseases, the majority of studies relate to viral infections. However, the mechanism of viral infection-mediated RNA splicing alterations differs from the one reported in this study.

Loss of hsr1 in *Mtb* resulted in a compromised growth of bacteria inside human macrophages or in mouse lungs. This observation makes *Mtb* hsr1 a novel virulence factor. While we cannot rule out additional mechanisms, however, since unlike the WT *Mtb*, the *Δhsr1* strains cannot alter host RNA splicing, we propose that losing the ability to alter host RNA splicing compromises the virulence of *Mtb*. This observation makes host RNA splicing regulators and the mycobacterial factors regulating that as a new set of host- and pathogen-directed therapy candidates against TB, respectively.

## Acknowledgements

The work was supported by DBT-Wellcome Trust India Alliance Senior Fellowship to DK (IA/S/17/1/503071). The work was also partly supported by grants from SERB (EMR/2016/005296 to DK) and the Department of Biotechnology, Govt of India (BT/IC-06/003/91-Flagship program to DK).

## Author Contributions

Original conceptualization: DK; Experiments: KC, NP, YK, DD, NA, PS, BM, HK, LS, LM; Investigation: KC; Analysis and Visualization: KC, MD, DK; Resources: VN, AS, VA, PD; Supervision: DK; Writing-original draft: DK, KC, MD, NP; Writing—review & editing: VN, AS, PD, DK

## Supplementary information for the manuscript

### Ethics statement

Studies on human samples were approved by the IEC of AIIMS, New Delhi (Ref no. IEC-304/02-06-2017). For animal experiments, approval was obtained from the Institutional Animal Ethics Committee, ICGEB, New Delhi (ICGEB/IAEC/22092022/CI-28).

### Biological materials

Tissue sections from Intestinal TB biopsy samples used in the study were obtained from the Department of Pathology, AIIMS, New Delhi. Access to these materials is subject to institutional guidelines.

### Clinical diagnosis of ITB subjects

The diagnoses depend on clinical features of nocturnal fever, weight loss, diarrhoea, and radiological suspicion of short-segment bowel thickenings, narrowed ileocaecal region, significant mesenteric lymphadenopathy, and/or necrosis. In such cases, an intestinal biopsy is taken, or a segment of the intestine is resected if there is a feature of intestinal obstruction due to fibrosis. All our cases underwent intestinal resection. Histological examination is critical in such cases, and demonstration of large, confluent, multiple, necrotic granulomas with multinucleated giant cells in deeper layers of the intestine are suggestive of ITB, as we have demonstrated in our cases. ZN stain has a very low sensitivity, varying from 5-10%. Such cases are confirmed by either the Gene-Xpert test, Mycobacterial culture, or where such tests are not possible, by ATT trial followed by re-evaluation at two months and six months. All our cases underwent either a *Mycobacterial* culture or a gene-Xpert test.

### Antibodies and reagents

The antibodies used in the immunofluorescence experiments were as follows: SNRPD2 (#NBP2-95236), SNRPF(#NBP1-57463), LSm7 (#NBP1-92084), Gemin3 (#NBP2-52454) and Gemin8(#NBP1-83925) from Novus biologicals.DDX23(#A300-693A) from Bethyl Laboratories were used. PKH26 red and PKH67 green fluorescent labelling dyes were used from Sigma-Aldrich. The spliceosome complex pull-down was done using an anti-SNRPB antibody (Y-12, MA513449). The antibodies used in immune-blotting experiments were as tubulin (Abcam plc; ab6046), histone (Abcam plc; ab18521), IRDye 800CW Goat anti-mouse IgG secondary antibody (LI-COR Biosciences)and IRDye 680RD goat anti-rabbit IgG secondary antibody(LI-COR Biosciences). The co-immunoprecipitation was done using EZView Red Anti-HA affinity gel (E6779), Anti-FLAG M2 Affinity gel (A2220) and immunoblots were detected using Anti-HA antibody, Mouse monoclonal (H3663) and Monoclonal Anti-FLAG M2 antibody mouse ( F1804) from Sigma-Aldrich.

### Cell culture

#### Mammalian cells

THP1 monocytes (ATCC) were maintained in RPMI 1640 media (Cell Clone), and HEK293T cells (ATCC) were maintained in DMEM media (Cell Clone). Cells were grown at 37°C with a 5% CO2 atmosphere. All the media was supplemented with 10% fetal bovine serum. HEK293T cells were transfected with plasmids at a 30-50% confluency using jetPRIME buffer (Polyplus #712-60) as per manufacturer protocol. The siRNA was transfected using the Dharmafect reagent. HEK293T constitutively expressing PRINT plasmid was selected in the presence of blasticidin (Gibco) 20 μg/ml concentration.

#### Bacterial Cells

*M. tuberculosis H37Rv*, *M. smegmatis* mc^2^155 and mutant strains were grown at 37°C to log phase in Middlebrook 7H9 media with 0.05% Tween80, 10% ADC Enrichment, and 0.2% glycerol. Plasmids were selected with 50 mg/ml kanamycin or hygromycin depending upon the resistance marker.

### Infection

THP-1 cells were activated into macrophages by treatment with 32 nM PMA (Sigma) for 24 hours, followed by another 24 hours for recovery. The THP-1-derived macrophages were infected with *Mtb* strains at an MOI of 10 for 4 hours, followed by 2 hours of amikacin (200μg/μl) treatment. *M. smegmatis* infection was performed at an MOI of 10 for 1 hour, followed by 2 hours of amikacin treatment. Amikacin treatment clears all the extracellular bacteria, which are washed following the treatment, and this time point is considered as 0 hours of infection throughout the manuscript.

### RNA isolation and qPCR

RNA was isolated at indicated time points using MDI mammalian Total RNA isolation kit (MTRK250) as per the manufacturer’s protocol. The RNA was quantified using a nanodrop spectrophotometer. 1000ng RNA was used to make cDNA using the Iscript cDNA synthesis kit (BioRad #170-8891), and 2 μl of 1:5 diluted cDNA was used to set qPCR on the BioRad CFX384 well plate qPCR machine. Primers were custom synthesized from Sigma Aldrich Chemicals Ltd. A list of primers used in this study is provided in the table S7.

### Transformation of yeast cell

The yeast competent cells were prepared by harvesting yeast cells by centrifugation at 400 rpm for 5 min at 25 °C. The supernatant was discarded, and the pellet was washed and resuspended in filter-sterilized SORB buffer (100 mM Lithium acetate, 100 mM Tris HCl, pH 8.0, 1 mM EDTA, pH 8.0, 1 M sorbitol pH 8.0) to which denatured salmon sperm DNA was added (10 mg/ml). The bait (pGBDKT7 vector) plasmids were mixed gently with the Y187 competent cells. Then PEG mixture was added and incubated at 30 °C for 30 min, 42 °C for 30 min and 4 °C for 30 min in a thermocycler machine. Tubes were taken out and plated on 15 cm diameter SC-TRP plates (Cat No. Clonetech 630413). Plates were incubated at 30°C for 2 to 7 days. A similar protocol was used for the prey (pGADT7 vector) plasmid in the AH109 yeast strain on SC-Leu medium (Cat No. Clonetech 630414).

### Y2H assay between bait and prey libraries

The bait library was prepared by plating the Y187 yeast strain (bait) transformed with Mtb proteins in 384 well patterns on the Singer Robot Plate. The AH109 (prey) yeast strain transformed with host spliceosome protein was grown in SC-leu broth medium and equally plated on a robotic compatible square plate. Using 384 pinned robotic pad bait libraries were patched on another YAPD plate. Each plate was then patched with individual cells containing prey plasmid and mixed well using the robotic system. This step was performed 4 times as four independent mating steps, thereby making it a total of 1536 spots per plate. The plate containing mixed colonies was kept for mating of both strains at 30 °C for 36 hours. After 36 hours these colonies were selected on SC-Trp-Leu (Cat No. Clonetech 630417) plates for the selection of those cells which contain both plasmids (bait as well as prey). After 72 hours these plates were replicated on SC-Trp-Leu-His (3 drops-out, Clonejet 630419) and SC-Trp-Leu-His-Ade (4 drop-outs, Clonetech 630428) for the screening of interaction between the protein which was fused with bait and prey respectively. The colonies that appear on both 3-DO and 4-DO plates represented very specific and strong interactions between the bait and prey proteins.

### Co-immunoprecipitation and Immunoblotting

The cells were lysed using a non-denaturing IP lysis Buffer (20 mM Tris-Cl, 137 mM NaCl, 2 mM EDTA, 10% glycerol, 1% NP-40, pH 8) and centrifuged at 14000 rpm for 10 min at 4 °C. The supernatant was precleared using IgG beads, and 300 μg of protein was used for IP. The IP was performed using FLAG-tagged beads (Sigma) and HA-tagged beads (Sigma) at 4 °C for 4 hours, followed by three washes with IP buffer. The beads were then boiled in 2X SDS loading dye. The protein samples were subjected to SDS PAGE followed by transfer on nitrocellulose membrane (BioRad) using BioRad Turbo transfer blot. The blocking was performed by Odessey blocking buffer (LI-COR Biosciences) in 1:1 dilution with 1X PBS at room temperature. The blots were immunoblotted with primary antibodies, followed by IR-tagged secondary antibodies (LI-COR Biosciences). Blot imaging was done using Odessey IR Imaging System (LI-COR Biosciences).

### Immunofluorescence

For immunofluorescence experiments, ∼0.2x 10^6^ cells were plated on 12 mm coverslips per well in a 24-well plate. Cells were infected with m-cherry expressing *Mtb*, and other strains were labelled with PKH67 fluorescent dye (Sigma). The cells were fixed using 4% v/v paraformaldehyde for 20 min at room temperature for indicated time points. The cells were incubated with blocking buffer (0.02% Triton-X100, 3% w/v BSA in 1XPBS) for 1 hour. The cells were washed thrice with 1X PBST and incubated with primary antibodies for 3 hours. The cells were washed thrice with 1XPBST and incubated with secondary Alexa Flour (488, 555, 590 or 670nm) tagged antibodies (Invitrogen). The nucleus was stained using DAPI (300 nM) for 10 min. The cells were mounted on slides with Pro-Long Gold antifade reagents (Life Technologies). All the incubation and staining were done at room temperature. Images were acquired by NIS-Elements software using the Nikon A1R laser scanning confocal microscope equipped with a Nikon Plan Apo VC ×20, NA 0.75, and Plan Apo VC ×100 oil, NA 1.40 objectives were used. Serial confocal sections, 0.5 µm thick, were acquired with a z-stack spanning 10–15 µm to form a composite image. Images were analysed using Imaris 9.6.6 imaging software using several tools like deconvolution, 3D-reconstruction and shortest distance calculation amongst others, NIS-Elements and image J software.

### Fluorescent reporter splicing assay using flow cytometry

For Flow cytometry experiments, ∼0.5 x 10^6^ HEK293T cells constitutively expressing pRINT vector (Addgene plasmid # 24218) were plated on 12 well plates. These were co-transfected with 800ng of modified pGINT vector (Addgene plasmid # 24217) with exon skipping cassette and *Mtb* protein-expressing vectors at a ratio of 1:1 using jetPRIME buffer. The cells were fixed using 4% v/v paraformaldehyde in 1X PBS for 20 minutes at room temperature. Then, the cells were scraped and resuspended in 1X PBS. The data were acquired using BD FACSDiva acquisition software in BD FACS Fortussa cytometer (BD).

### Spliceosome complex Co-IP

For the pulldown experiment, ∼200 million THP-1 derived macrophages were infected with H37Rv at an MOI of 10 and samples were processed 48-hour post-infection. The nuclear and cytoplasmic extract was prepared using the Active Motif Nuclear Complex Co-IP kit (catalogue No.54001) per the manufacturer’s guidelines. The nuclear extract and cytoplasmic extract were passed through a 0.2-micron filter. 500 µg of nuclear extract and cytoplasmic extracts were used for pulldown using 10 µg Y-12 antibody (MA5-13449, Thermo Fisher) or IgG-control (Sigma) antibody for each condition. The Y-12 antibody was crosslinked using DMP (20 mM) to Protein A-Sepharose beads for 1 hour (beads were washed prior with 0.2 M sodium borate, pH 9.0) followed by 0.2 M ethanolamine incubation for 1 hour at room temperature. The Nuclear and Cytosolic extracts were then incubated with crosslinked beads overnight at 4 °C. The beads were washed thrice with wash buffer (0.025 M Tris, 0.15 M NaCl, pH 7.2). The protein was finally eluted in 50 µl of an elution buffer containing primary amine pH 2.8. To neutralize the low pH of the elution buffer, 1 M Tris (pH 9.5) was added to the collection tube.

### Mass Spectrometric Data analysis for immune-precipitated spliceosome complex

*Sample Preparation*-An equal volume of eluted protein sample (50ul) was used for digestion and reduced with 5 mM TCEP and further alkylated with 50 mM iodoacetamide and then digested with Trypsin (1:50, Trypsin/lysate ratio) for 16 h at 37 °C. Digests were cleaned using a C18 silica cartridge to remove the salt and dried using a speed vac. The dried pellet was resuspended in buffer A (2% acetonitrile, 0.1% formic acid). An equal amount of peptide (1ug) was injected into the analytical column. *Mass Spectrometric Analysis of Peptide Mixtures*-All the experiments were performed using EASY-nLC 1000 system (Thermo Fisher Scientific) coupled to Thermo Fisher-QExactive plus equipped with nano electrospray ion source. 1ug peptide was loaded on C18 column 50 cm, 3.0μm Easy-spray column (Thermo Fisher Scientific) at column heating temp 40 C. Peptides were eluted with a 0–40% gradient of buffer B (80% acetonitrile, 0.1% formic acid) at a flow rate of 300 nl/min) and injected for MS analysis. LC gradients were run for 90 minutes MS1 spectra were acquired in the Orbitrap (Max IT = 50ms, AGC target = 3e6; R=70K, mass range = 350-1750; Profile data). Dynamic exclusion was employed for 30s excluding all charge states for a given precursor. MS2 spectra were collected for top 15 peptides. MS2 (Max IT= 100ms, R= 17.5K, AGC target 1e5). *Data Processing-* All samples were processed and RAW files generated were analysed with Proteome Discoverer (v2.2) against the Uniprot Human and Mtb database. For dual Sequest and Amanda search, the precursor and fragment mass tolerances were set at 10 ppm and 0.5 Da, respectively. The protease used to generate peptides, i.e. enzyme specificity was set for trypsin/P (cleavage at the C terminus of “K/R: unless followed by “P”). Carbamidomethyl on cysteine as fixed modification and oxidation of methionine, N-terminal acetylation were considered as variable modifications for database search. Both peptide spectrum match and protein false discovery rate were set to 0.01 FDR. Only statistically significant proteins (Using ANNOVA p-value cut off 0.05) that were found in all three biological replicates across and in all experimental groups were used to generate heatmap and for further analysis.

### RNA FISH

RNA-fluorescence *in situ* hybridization (RNA FISH) was performed using previously described snRNA probes ^1^. The sequences of the oligonucleotide used for FISH were as follows: U5 snRNA, 5′-CTC TCC ACG GAA ATC TTT AGT AAA AGG CGA AAG ATT TAT ACG ATT TGA AGA G-3′; The U5 snRNA probes were labelled with Alexa Flour 647 (Invitrogen) at their 5’ end. Next, the cells were fixed with 4% v/v paraformaldehyde for 20 min at room temperature. Next, cells were permeabilized and incubated with RNA-FISH probe for three hours at 40 °C by using the ViewRNA™ ISH Cell Assay Kit system (Thermo Fischer). Subsequently, cells were immune-stained and mounted using a Pro-Long gold antifade (Life technologies).

### Bacterial transformations

For the generation of *M. smegmatis* mutants expressing *H37Rv* proteins, 50 ml of an early exponential phase culture of *M. smegmatis* mc^2^155 was washed twice and concentrated ice-cold 10% ultrapure glycerol (Invitrogen). Concentrated cells were electroporated with approximately 10 μg of pNit plasmid. Electroporation was carried out at 2500 V, 25 μF, and 1000 Ω in the Biorad gene pulsar. Electroporated cells were allowed to recover at 37 °C in 7H9 Middlebrook broth and plated on 7H11 supplemented with 10% OADC and kanamycin (25 μg/ml; Sigma).

### *Mtb* survival assay

For CFU experiments, ∼0.01 million THP-1 macrophages were seeded in a 96-well plate and *Mtb* infection was done at an MOI of 10. The lysis was done by incubating with 0.06% SDS for 5 min at indicated time points. The lysed cells were diluted to 1:10 and 1:50 in 7H9 media and 10 µl of the dilutions were plated in a drop-wise fashion on a 7H11 plate supplemented with 10% OADC. The plates were incubated for approximately around 3 weeks for colony enumeration at 37 °C.

### Expression and purification of recombinant proteins

The Mtb proteins were cloned in the pEZY19 vector and transformed into E. coli BL21(DE3) competent. Protein expression was induced with 0.1 mM isopropyl b-D-1-thiogalactopyranoside (IPTG) for 16-19 hours. Collected frozen cells were lysed in a lysis Buffer (50 mM Tris-HCl pH. 8.0 containing 300 mM NaCl,10% Glycerol, 1mm DTT, 1mm PMSF,1mm BHCL,0.1% Triton X-100, 10mM imidazole and lysozyme 1mg/ml) followed by sonication (0.5 s pulse and 0.9 s rest) at 2 min intervals to prevent overheating. After centrifugation, the proteins were purified from the supernatant using nickel-nitrilotriacetic acid (Ni-NTA) affinity chromatography (Cat. No.-30410, Sigma). The recombinant proteins were eluted from Ni-NTA resins by elution buffer (50 mM Tris-HCl pH. 8.0 and 300 mM NaCl) containing 300mm Imidazole. The purified protein was concentrated using 10Kda Centricon (ACS501024, Millipore) subjected to gel permeation chromatography on GE Healthcare Superdex 75 Increase 10/300 GL where the monomeric fraction of the protein was eluted. The final protein was stored in 50 mM Tris-HCl pH. 8.0 containing 300 mM NaCl,10% Glycerol buffer at −80°C.

### BALB/c aerosol challenge

All mice experiments were carried out in the Tuberculosis Aerosol Challenge Facility (TACF, ICGEB, New Delhi, India). BALB/c mice have been housed in individually ventilated cages contained within the biosafety level 3 enclosure. The animal holding area of the BSL3 lab is maintained at 20–25 °C, 30–60% humidity and 12–12 h of the light–dark cycle. An aerosol challenge of ∼100 CFUs was given to the animals in a Wisconsin–Madison chamber according to the standardized protocol in the TACF facility. Two animals were selected randomly and humanely euthanized 24 h post aerosol challenge to check for infection establishment. The lungs and spleen tissues were harvested and homogenized to enumerate CFU. Tissue lysates were serially diluted and plated on Petri plates containing Middlebrook 7H11 agar (Difco) supplemented with 10% OADC (Becton, Dickinson) and 0.5% glycerol.

### RNA-Seq analysis

Reads were quality filtered using Trimmomatic (v.0.39, LEADING:10 TRAILING:10 SLIDING WINDOW:4:15 MINLEN:75) ^2^ and were assessed for quality using FastQC (v.10.1) ^3^ and visualised with MultiQC (v.1.9)^4^. Salmon (v.1.8, –numGibbsSamples 30) ^5^ was used to quantify quality filtered reads against the human genome (GRCh38, primary assembly) with Gencode annotation (v.40) ^6^. Profiles were imported using tximeta (v.1.10) ^7^ and ^8^ (v.1.22) and filtered to retain all transcripts from only protein-coding genes. Statistical analysis was done using swish methodology from fishpond (v.1.8) ^9^ and significantly differentially expressed transcripts (DETs) were defined based on *qvalue* ≤ 0.1. Furthermore, Isoform usage was analysed using IsoformSwitchAnalyzeR (v.1.18) ^10^ with reduceToSwitchingGenes = FALSE, reduceFurtherToGenesWithConsequencePotential = FALSE, No dIFcutoff and Isoform fraction cuffoff. Heatmap of the DETs was generated with ^11^ using mean z-scaled normalised scaled TPM values and clustered based on Euclidean distance and Ward.D2 methodology. Clusters were defined manually. Enrichment analysis was done separately on overexpressed and repressed DETs from each pairwise comparisons using enrichR (v.3.0) ^12^ with KEGG Human 2021 and Gene Ontology (GO) ^13^ (Biological Process (BP), Cellular Component (CC) and Molecular Function (MF)) 2021 database figures were generated using ggplot2 (v.3.3.5)^14^. Analyses were done in R statistical software environment (v.4.1) ^15^.

### Generation of Rv1435c mutant

*Rv1435c* mutant was generated using recombineering methodology which requires an Allelic exchange substrate ^16^. To construct that, 836bp upstream (LHS) and 1Kb downstream (RHS) regions of the *Rv1435c* locus (retaining ∼170 bp of the gene at both ends) were amplified from *Mtb* (*H37Rv* genome) with the help of primers having appropriate restriction endonuclease sites (LHS-DraIII; RHS-SfiI). The flanks were then digested and ligated with other two fragments-Hygromycin resistant cassette and *oriE*+cosλ with compatible ends to create a four-piece AES. The latter was then digested with SnaBI sites to release the linearised fragment consisting of LHS-*Hyg^r^*-RHS and electroporated into recombineering proficient *Rv::pNitET* (Kan^r^) strain that overexpresses RecET recombinases upon over-expression using Isovaleronitrile. Mutant colonies were analysed using different primer sets to confirm legitimate recombination at the required locus and cured of the pNitET plasmid by negative selection on 2% sucrose-containing 7H11-hyg^r^ agar plates.

Primers used:

*Rv1435c* 5’ Flank forward primer with DraIII & SnaBI site:

5’CACCTTTTCACAAAGTGTACGTACGTGGTGCACGACGCATTGGA 3’

*Rv1435c* 5’ Flank reverse primer with DraIII site:

5’ TTTTTTTTCACTTCGTGCGCCGGCGGCTACCGGCGCGT 3’

*Rv1435c* 3’ Flank forward primer with SfiI site:

5’ CACCTTTTGGCCTAGATGGCCGGGCCGGTCCTGGCTGGTGCT 3’

*rv1435c* 3’ Flank reverse primer with SfiI site and SnaB1 site:

5’ TTTTTTTTGGCCACTTAGGCCTACGTAGGGATCGCTGGGAGCGTGGTT 3’

### Generation of Rv1435c complementation construct pSCW-F-Rv1435c

*Rv1435c* was cloned along with its native promoter (500bp upstream of the gene) between ScaI and HindIII sites using an integrative vector, pSCW-F (C-terminal 3X FLAG tag), which facilitates the integration of the gene at Tweety attP sites in the *Mtb* genome.

Primers used:

FP *Rv1435c* (with UPSTREAM 500bp) ScaI

5’ FP : 5’ CACCAGTACTATGCGCCAGCGTGCTGTTG 3’

RP *Rv1435c* HindIII, w/o stop codon

5’TTTTAAGCTTGCGCGCACCTGCGGGC 3’

#### Immunohistochemical staining

The paraffin-embedded specimens were cut into 4–5 µm-thick sections, deparaffinized in xylene for 5–10 min, and dehydrated through a series of graded alcohols (100%, 100%, 95% and 80%). Tissue sections were boiled at 100°C for 30 min in 10mM citrate buffer (pH 6.0), followed by cooling and treatment with blocking solution (100% methanol containing 0.3% H_2_O_2_) for 30 min to block any endogenous peroxidase activity. They were incubated with SNRPF or SNRPD2 primary antibody (1:600, pH 9) overnight at 4°C, followed by incubation with secondary antibody (Ultra trek anti polyvalent kit with HRP conjugated secondary kit, Scy Tek Laboratories, Logan, USA) for 30 mins at room temperature. The slides were thereafter stained with 3,3′-Diaminobenzidine (DAB). Finally, the sections were counterstained with Hematoxylin and Eosin and observed under a microscope.

#### Interpretation of IHC results

SNRPF staining in >10% of cells was considered positive. Both cytoplasmic and nuclear staining were looked for. The spatial distribution of the stained sections, in terms of granulomas areas and areas away from granulomas, were noted down to identify polarization of staining, if any.

### Legends to Figures S1-S9

**Figure S1:**
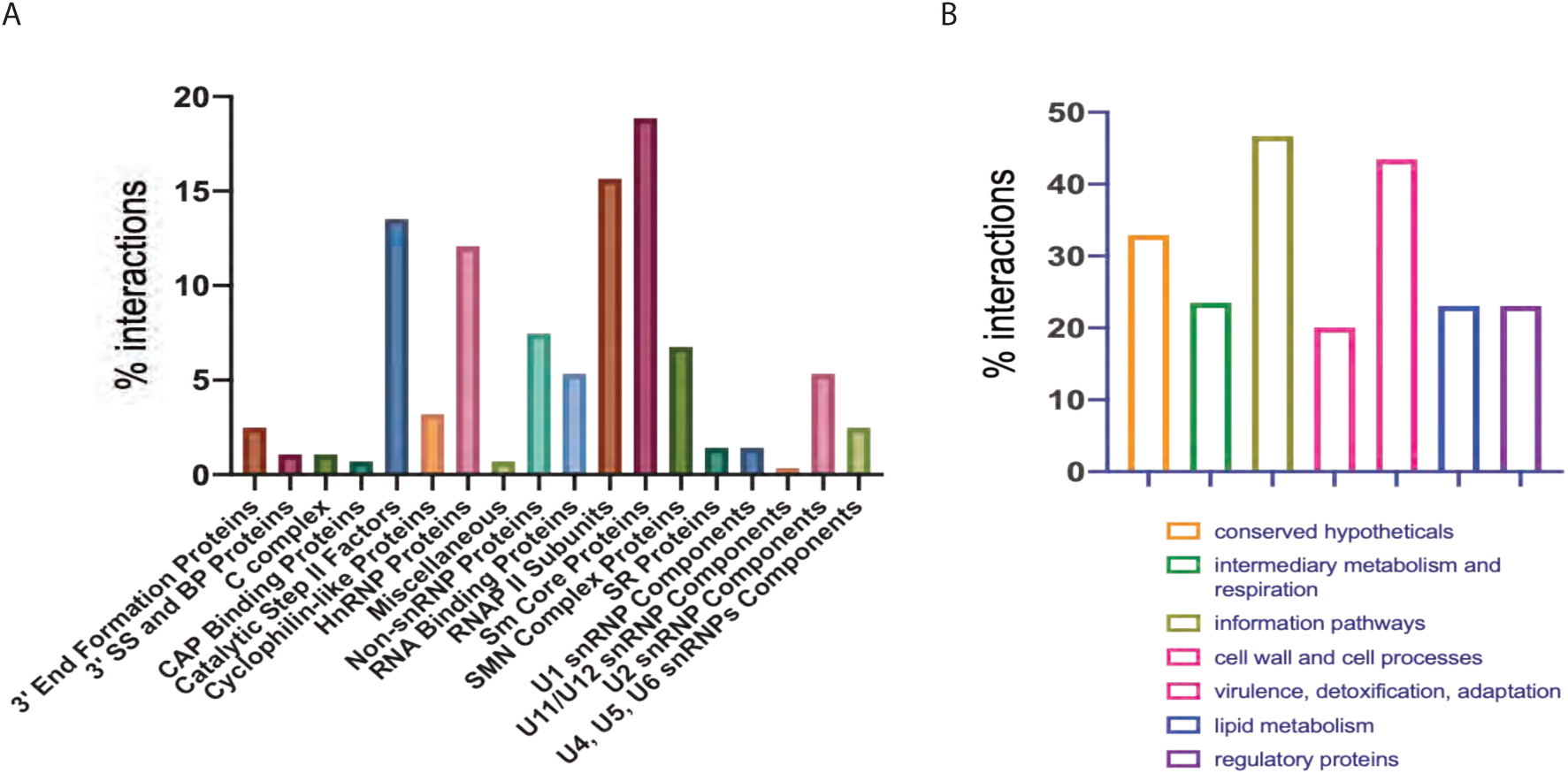
Mycobacterial secretory proteins and host spliceosome proteins interactome. **A)** The percentage of the interaction shown by spliceosome-related proteins belonging to different spliceosome protein categories **B)** The percentage of the interacting Mtb proteins belonging to the different functional classifications.

**Figure S2:**
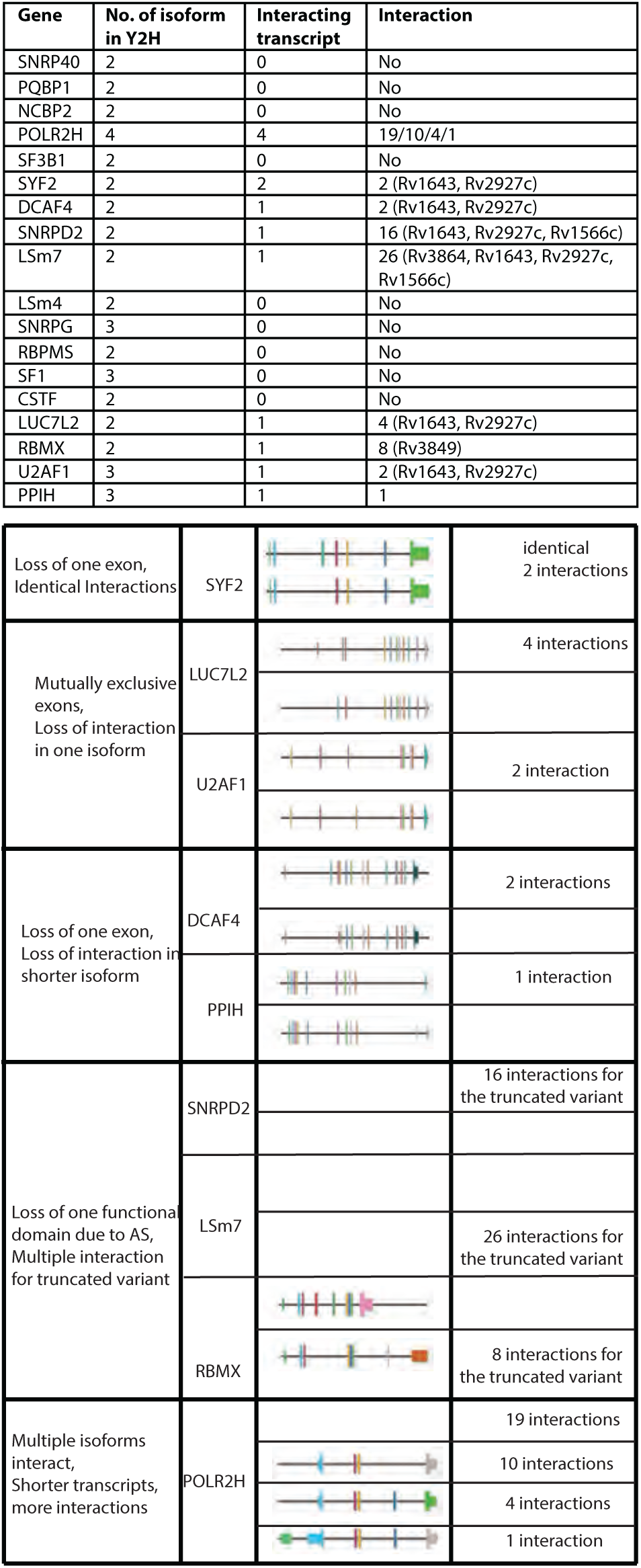
Interaction of Mtb proteins with the splice variants of human proteins. A) A descriptive list of the human spliceosome proteins, represented by more than one isoform in the AD library, along with their interactions with the *Mtb* protein, is presented in the table. B) The table shows the distinct spliced mRNA isoforms that are classified as loss of exon, mutually exclusive exon and loss of the functional domain. The isoforms of SYF2, differing by only one exon, show interaction with the same *Mtb* proteins, suggesting no contribution of alternatively spliced exons towards interaction. Only one of the isoforms of genes which show exon skipping, like DCAF4 and PPIH, or mutually exclusive exons like U2AF1 and LUC7L2, showed interaction with any *Mtb* proteins, suggesting the involvement of alternative exons in the interactions. The truncated isoforms of SNRPD2, LSm7 and RBMX show a loss of functional domains and multiple interactions with Mtb proteins, while the longer isoforms for these genes did not show any interactions. Similarly, the truncated isoforms of POLR2H show more interaction.

**Figure S3:**
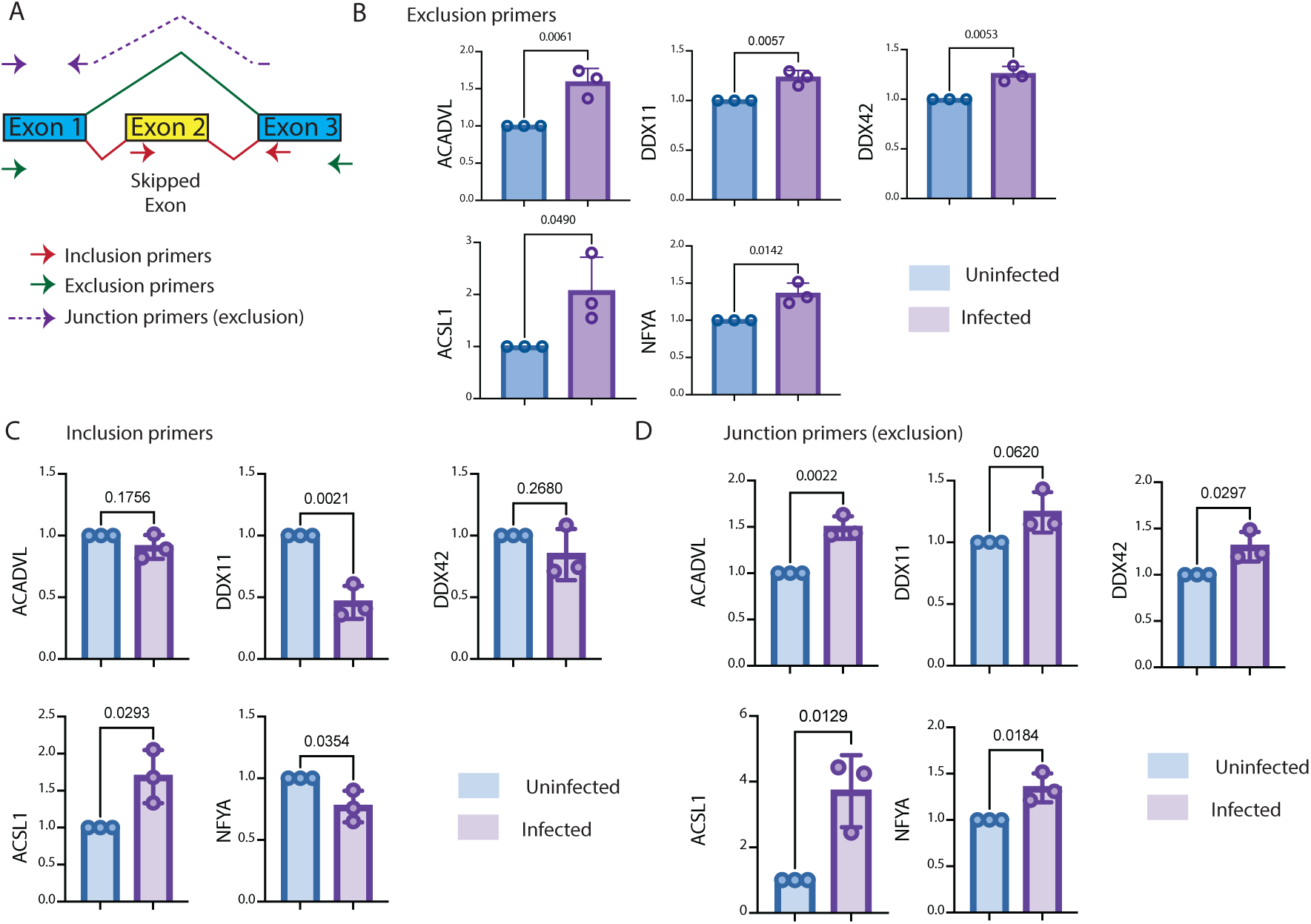
Altered RNA splicing in *Mtb*-infected THP-1 macrophages. **A)** The schematic shows the primer design strategy for exon-inclusion, exon-exclusion and exon-junction primers. **B)** Expression of five target genes (as indicated) in THP-1 macrophages upon infection with H37Rv using exon exclusion primers at 48 hpi. Data from three independent biological experiments. **C)** Expression of five target genes (as indicated) in THP-1 macrophages upon infection with H37Rv using exon inclusion primers at 48 hpi. Data from three independent biological experiments. **D)** Expression of five target genes (as indicated) in THP-1 macrophages upon infection with H37Rv using exon junction primers at 48 hpi. Data from three independent biological experiments.

**Figure S4:**
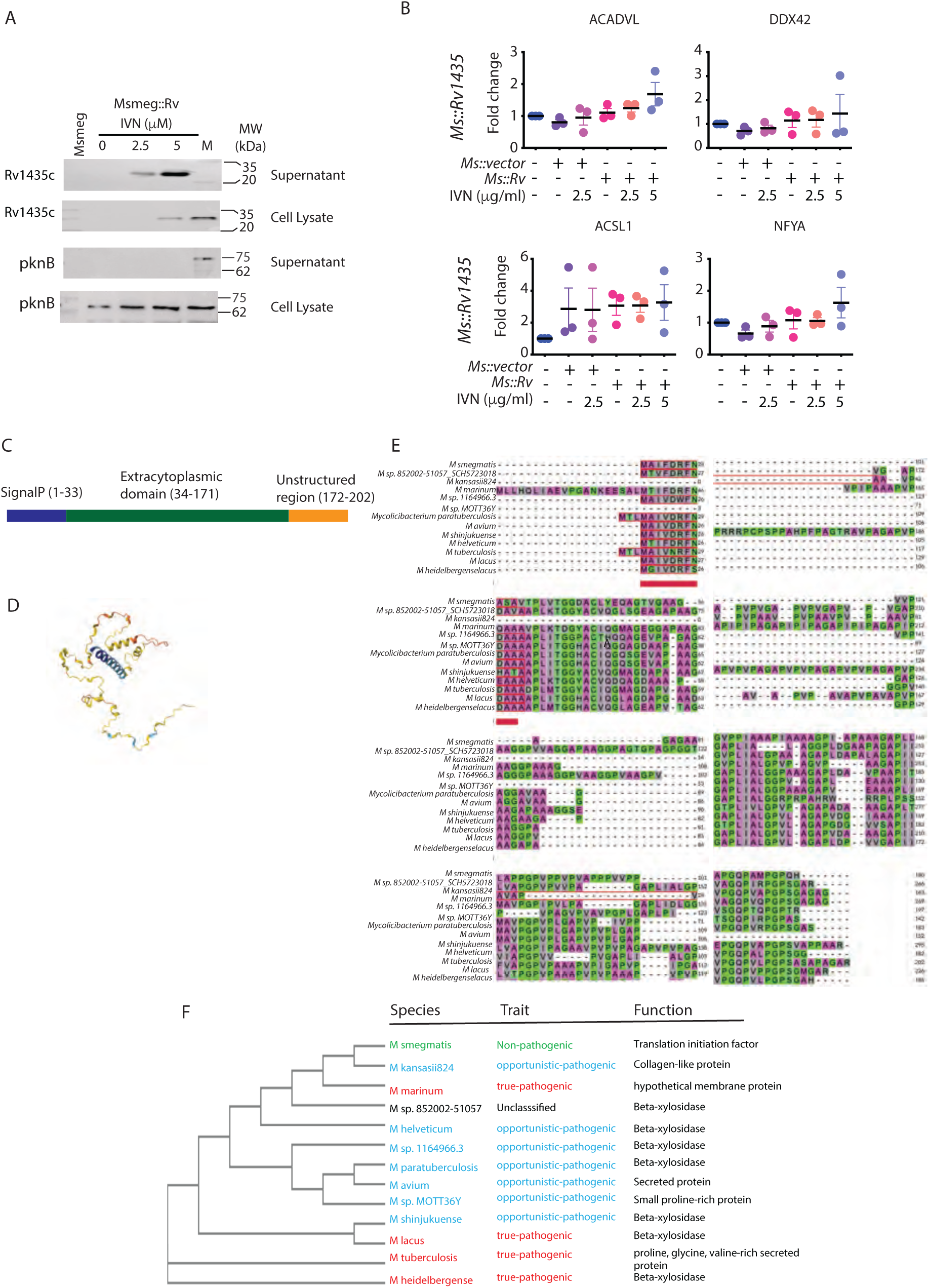
Rv1435 secretion from the modified *M.smegmatis* strains. **A)** The protein expression in modified *M. smegmatis* strains expressing Rv1435c upon induction with 2.5µM and 5 µM IVN was checked by immunoblotting in whole cell lysate and culture supernatant (SN). As control, the modified *M. smegmatis* strains expressing pknB were also induced with IVN and its expression in the supernatant or whole cell lysate was measured by immunoblotting. **B)** Isoform-specific qPCR to assess the impact of Rv1435c expressing *Msmeg* infection in THP-1 macrophages on exon skipping event in *ACADVL*, *DDX42*, *ACSL1* and *NFYA*; N=3, indicating p values from One-way ANOVA. **C)** Domain organization of Rv1435c depicting signalP, extracytoplasmic and disordered regions. **D)** Predicted structure of Rv1435c from AlphaFold Protein Structure Database **E)** Multiple sequence alignment of Rv1435c protein in different mycobacterial species using CLUSTAL Omega. The red bar indicates the conserved sequence among different species. **F)** A phylogenetic tree corresponding to the multiple sequence alignment is shown in S6C.

**Figure S5:**
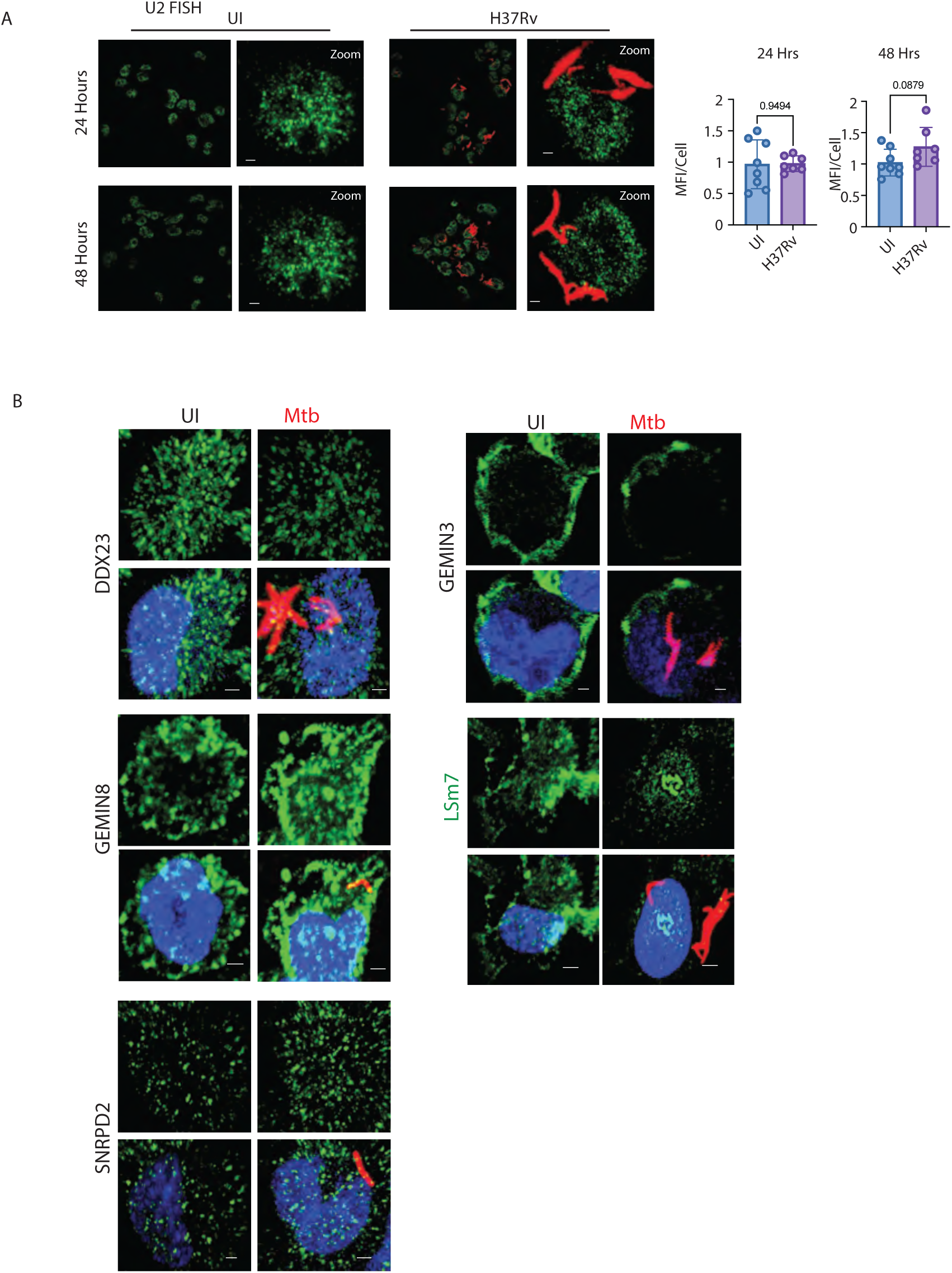
Dysregulation of splicing proteins upon Mtb infection. A) Representative images for RNA FISH against U2 snRNA in THP-1 macrophages upon *Mtb* (mCherry-H37Rv) infection at 24- and 48-hrs post-infection. The U5 snRNA probe was labelled with Alexa 488 (green). Plots on the right show quantification of U2 snRNA FISH MFI in uninfected and *Mtb*-infected macrophages at 24 ad 48 hpi. B) Representative image for staining of DDX23, GEMIN3, GEMIN8, LSm7 and SNRPD2 in uninfected and H37Rv infected THP-1 macrophages 48 h post-infection.

**Figure S6:**
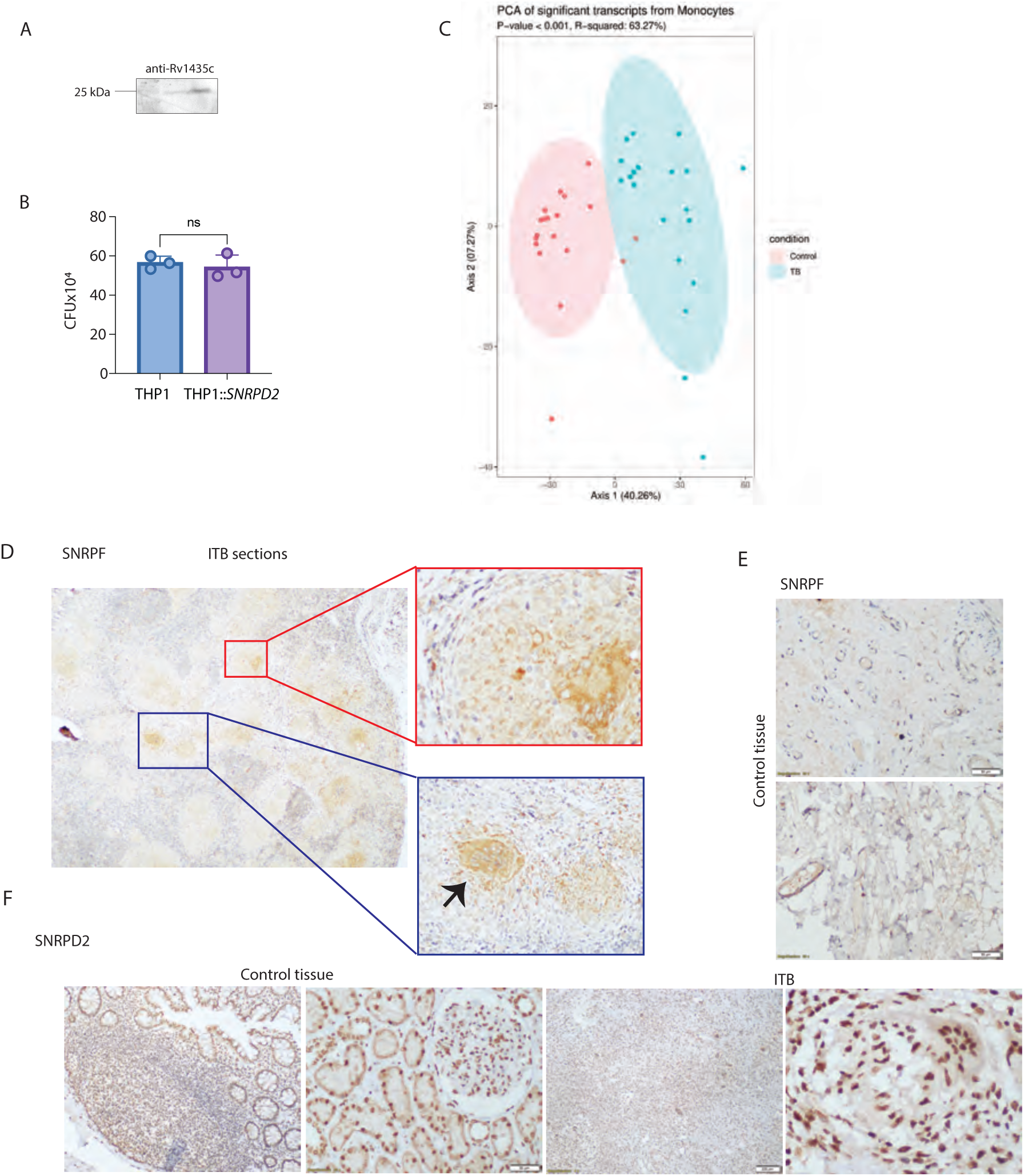
Regulation of host alternative splicing as a virulence mechanism. A) Western blot showing the specificity of anti-sera raised against Rv1435c. B) The plot represents the bacterial CFU in WT THP-1 and THP-1::SNRPD2 at 48hr post-infection. C) PCA analysis of significant transcripts (splice isoforms) identified by RNA-Seq in monocytes of Human TB patients. D) Human intestinal tuberculosis granulomatous lesions were stained with SNRPF. Intense cytosolic staining in multinucleated giant cells (black arrow) is shown. E) Representative image for IHC staining of SNRPF in gut sections from healthy controls. F) Representative image for IHC staining of SNRPD2 in gut sections from healthy controls and in intestinal tuberculosis granulomatous lesions from diseased individuals.

### Supplemental Tables

**Table S1**: Spliceosome proteins identified in Y-12 pulldown MS analysis

**Table S2**: List of Mtb genes in the bait library (BD library) and human splicing factor coding transcripts in the prey library (AD library).

**Table S3**: List of genes included in the BD library as negative controls

**Table S4**: Yeast-2-hybrid interaction matrix for AD and BD proteins

**Table S5**: Two-column file for the interaction network shown in Fig 2B.

**Table S6**: RNAseq data shown in Fig. 8A.

**Table S7**: List of primers

## References

1 Augenstreich, J. & Briken, V. Host Cell Targets of Released Lipid and Secreted Protein Effectors of Mycobacterium tuberculosis. Front Cell Infect Microbiol 10, 595029 (2020). 10.3389/fcimb.2020.595029

2 Chandra, P. et al. Mycobacterium tuberculosis Inhibits RAB7 Recruitment to Selectively Modulate Autophagy Flux in Macrophages. Sci Rep 5, 16320 (2015). 10.1038/srep16320

3 de Martino, M., Lodi, L., Galli, L. & Chiappini, E. Immune Response to Mycobacterium tuberculosis: A Narrative Review. Front Pediatr 7, 350 (2019). 10.3389/fped.2019.00350

4 Ernst, J. D. The immunological life cycle of tuberculosis. Nat Rev Immunol 12, 581–591 (2012). 10.1038/nri3259

5 Flynn, J. L. & Chan, J. Immunology of tuberculosis. Annu Rev Immunol 19, 93–129 (2001). 10.1146/annurev.immunol.19.1.93

6 Kumar, D. & Rao, K. V. Regulation between survival, persistence, and elimination of intracellular mycobacteria: a nested equilibrium of delicate balances. Microbes Infect 13, 121–133 (2011). 10.1016/j.micinf.2010.10.009

7 Zhai, W., Wu, F., Zhang, Y., Fu, Y. & Liu, Z. The Immune Escape Mechanisms of Mycobacterium Tuberculosis. Int J Mol Sci 20 (2019). 10.3390/ijms20020340

8 Karim, A. F. et al. Express path analysis identifies a tyrosine kinase Src-centric network regulating divergent host responses to Mycobacterium tuberculosis infection. J Biol Chem 286, 40307–40319 (2011). 10.1074/jbc.M111.266239

9 Koul, A., Herget, T., Klebl, B. & Ullrich, A. Interplay between mycobacteria and host signalling pathways. Nat Rev Microbiol 2, 189–202 (2004). 10.1038/nrmicro840

10 Lavalett, L. et al. Alveolar macrophages from tuberculosis patients display an altered inflammatory gene expression profile. Tuberculosis (Edinb) 107, 156–167 (2017). 10.1016/j.tube.2017.08.012

11 Lesho, E. et al. Transcriptional responses of host peripheral blood cells to tuberculosis infection. Tuberculosis (Edinb) 91, 390–399 (2011). 10.1016/j.tube.2011.07.002

12 McGarvey, J. A., Wagner, D. & Bermudez, L. E. Differential gene expression in mononuclear phagocytes infected with pathogenic and non-pathogenic mycobacteria. Clin Exp Immunol 136, 490–500 (2004). 10.1111/j.1365-2249.2004.02490.x

13 Kalam, H., Fontana, M. F. & Kumar, D. Alternate splicing of transcripts shape macrophage response to Mycobacterium tuberculosis infection. PLoS Pathog 13, e1006236 (2017). 10.1371/journal.ppat.1006236

14 Kalam, H., Singh, K., Chauhan, K., Fontana, M. F. & Kumar, D. Alternate splicing of transcripts upon Mycobacterium tuberculosis infection impacts the expression of functional protein domains. IUBMB Life 70, 845–854 (2018). 10.1002/iub.1887

15 Chauhan, K., Kalam, H., Dutt, R. & Kumar, D. RNA Splicing: A New Paradigm in Host-Pathogen Interactions. J Mol Biol 431, 1565–1575 (2019). 10.1016/j.jmb.2019.03.001

16 Artemyeva-Isman, O. V. & Porter, A. C. G. U5 snRNA Interactions With Exons Ensure Splicing Precision. Front Genet 12, 676971 (2021). 10.3389/fgene.2021.676971

17 Lee, Y. & Rio, D. C. Mechanisms and Regulation of Alternative Pre-mRNA Splicing. Annu Rev Biochem 84, 291–323 (2015). 10.1146/annurev-biochem-060614-034316

18 Matera, A. G. & Wang, Z. A day in the life of the spliceosome. Nat Rev Mol Cell Biol 15, 108–121 (2014). 10.1038/nrm3742

19 Papasaikas, P., Tejedor, J. R., Vigevani, L. & Valcarcel, J. Functional splicing network reveals extensive regulatory potential of the core spliceosomal machinery. Mol Cell 57, 7–22 (2015). 10.1016/j.molcel.2014.10.030

20 Schlautmann, L. P. & Gehring, N. H. A Day in the Life of the Exon Junction Complex. Biomolecules 10 (2020). 10.3390/biom10060866

21 Wahl, M. C., Will, C. L. & Luhrmann, R. The spliceosome: design principles of a dynamic RNP machine. Cell 136, 701–718 (2009). 10.1016/j.cell.2009.02.009

22 Wilkinson, M. E., Charenton, C. & Nagai, K. RNA Splicing by the Spliceosome. Annu Rev Biochem 89, 359–388 (2020). 10.1146/annurev-biochem-091719-064225

23 Gubitz, A. K., Feng, W. & Dreyfuss, G. The SMN complex. Exp Cell Res 296, 51–56 (2004). 10.1016/j.yexcr.2004.03.022

24 Karijolich, J. & Yu, Y. T. Spliceosomal snRNA modifications and their function. RNA Biol 7, 192–204 (2010). 10.4161/rna.7.2.11207

25 Liu, J. L. & Gall, J. G. U bodies are cytoplasmic structures that contain uridine-rich small nuclear ribonucleoproteins and associate with P bodies. Proc Natl Acad Sci U S A 104, 11655–11659 (2007). 10.1073/pnas.0704977104

26 Patel, S. B. & Bellini, M. The assembly of a spliceosomal small nuclear ribonucleoprotein particle. Nucleic Acids Res 36, 6482–6493 (2008). 10.1093/nar/gkn658

27 Pellizzoni, L. Chaperoning ribonucleoprotein biogenesis in health and disease. EMBO Rep 8, 340–345 (2007). 10.1038/sj.embor.7400941

28 Shaw, D. J., Eggleton, P. & Young, P. J. Joining the dots: production, processing and targeting of U snRNP to nuclear bodies. Biochim Biophys Acta 1783, 2137–2144 (2008). 10.1016/j.bbamcr.2008.07.025

29 Yong, J., Kasim, M., Bachorik, J. L., Wan, L. & Dreyfuss, G. Gemin5 delivers snRNA precursors to the SMN complex for snRNP biogenesis. Mol Cell 38, 551–562 (2010). 10.1016/j.molcel.2010.03.014

30 Zhang, R. et al. Structure of a key intermediate of the SMN complex reveals Gemin2’s crucial function in snRNP assembly. Cell 146, 384–395 (2011). 10.1016/j.cell.2011.06.043

31 Zhang, Z. et al. SMN deficiency causes tissue-specific perturbations in the repertoire of snRNAs and widespread defects in splicing. Cell 133, 585–600 (2008). 10.1016/j.cell.2008.03.031

32 Strasser, A., Dickmanns, A., Luhrmann, R. & Ficner, R. Structural basis for m3G-cap-mediated nuclear import of spliceosomal UsnRNPs by snurportin1. EMBO J 24, 2235–2243 (2005). 10.1038/sj.emboj.7600701

33 Wang, Z. & Burge, C. B. Splicing regulation: from a parts list of regulatory elements to an integrated splicing code. RNA 14, 802–813 (2008). 10.1261/rna.876308

34 Cartegni, L., Hastings, M. L., Calarco, J. A., de Stanchina, E. & Krainer, A. R. Determinants of exon 7 splicing in the spinal muscular atrophy genes, SMN1 and SMN2. Am J Hum Genet 78, 63–77 (2006). 10.1086/498853

35 Pai, A. A. et al. Widespread Shortening of 3’ Untranslated Regions and Increased Exon Inclusion Are Evolutionarily Conserved Features of Innate Immune Responses to Infection. PLoS Genet 12, e1006338 (2016). 10.1371/journal.pgen.1006338

36 Penn, B. H. et al. An Mtb-Human Protein-Protein Interaction Map Identifies a Switch between Host Antiviral and Antibacterial Responses. Mol Cell 71, 637–648 e635 (2018). 10.1016/j.molcel.2018.07.010

37 Portal-Celhay, C. et al. Mycobacterium tuberculosis EsxH inhibits ESCRT-dependent CD4(+) T-cell activation. Nat Microbiol 2, 16232 (2016). 10.1038/nmicrobiol.2016.232

38 Sharma, G., Upadhyay, S., Srilalitha, M., Nandicoori, V. K. & Khosla, S. The interaction of mycobacterial protein Rv2966c with host chromatin is mediated through non-CpG methylation and histone H3/H4 binding. Nucleic Acids Res 43, 3922–3937 (2015). 10.1093/nar/gkv261

39 Yaseen, I., Kaur, P., Nandicoori, V. K. & Khosla, S. Mycobacteria modulate host epigenetic machinery by Rv1988 methylation of a non-tail arginine of histone H3. Nat Commun 6, 8922 (2015). 10.1038/ncomms9922

40 Klimesova, K. et al. TSSC4 is a component of U5 snRNP that promotes tri-snRNP formation. Nat Commun 12, 3646 (2021). 10.1038/s41467-021-23934-y

41 Neuenkirchen, N. et al. Reconstitution of the human U snRNP assembly machinery reveals stepwise Sm protein organization. EMBO J 34, 1925–1941 (2015). 10.15252/embj.201490350

42 Malakar, B. et al. Phosphorylation of CFP10 modulates Mycobacterium tuberculosis virulence. mBio 14, e0123223 (2023). 10.1128/mbio.01232-23

43 de Souza, G. A., Leversen, N. A., Malen, H. & Wiker, H. G. Bacterial proteins with cleaved or uncleaved signal peptides of the general secretory pathway. J Proteomics 75, 502–510 (2011). 10.1016/j.jprot.2011.08.016

44 Malen, H., Berven, F. S., Fladmark, K. E. & Wiker, H. G. Comprehensive analysis of exported proteins from Mycobacterium tuberculosis H37Rv. Proteomics 7, 1702–1718 (2007). 10.1002/pmic.200600853

45 Mehra, A. et al. Mycobacterium tuberculosis type VII secreted effector EsxH targets host ESCRT to impair trafficking. PLoS Pathog 9, e1003734 (2013). 10.1371/journal.ppat.1003734

46 Bonano, V. I., Oltean, S. & Garcia-Blanco, M. A. A protocol for imaging alternative splicing regulation in vivo using fluorescence reporters in transgenic mice. Nat Protoc 2, 2166–2181 (2007). 10.1038/nprot.2007.292

47 Luan, X. et al. High Immunogenicity of a T-Cell Epitope-Rich Recombinant Protein Rv1566c-444 From Mycobacterium tuberculosis in Immunized BALB/c Mice, Despite Its Low Diagnostic Sensitivity. Front Immunol 13, 824415 (2022). 10.3389/fimmu.2022.824415

48 Varadi, M. et al. AlphaFold Protein Structure Database in 2024: providing structure coverage for over 214 million protein sequences. Nucleic Acids Res 52, D368–D375 (2024). 10.1093/nar/gkad1011

49 van Kessel, J. C. & Hatfull, G. F. Recombineering in Mycobacterium tuberculosis. Nat Methods 4, 147–152 (2007). 10.1038/nmeth996

50 Massenet, S., Pellizzoni, L., Paushkin, S., Mattaj, I. W. & Dreyfuss, G. The SMN complex is associated with snRNPs throughout their cytoplasmic assembly pathway. Mol Cell Biol 22, 6533–6541 (2002). 10.1128/MCB.22.18.6533-6541.2002

51 Wood, K. A., Eadsforth, M. A., Newman, W. G. & O’Keefe, R. T. The Role of the U5 snRNP in Genetic Disorders and Cancer. Front Genet 12, 636620 (2021). 10.3389/fgene.2021.636620

52 Will, C. L. et al. A novel U2 and U11/U12 snRNP protein that associates with the pre-mRNA branch site. EMBO J 20, 4536–4546 (2001). 10.1093/emboj/20.16.4536

53 Becker, D. et al. Nuclear Pre-snRNA Export Is an Essential Quality Assurance Mechanism for Functional Spliceosomes. Cell Rep 27, 3199–3214 e3193 (2019). 10.1016/j.celrep.2019.05.031

54 Jia, Y., Mu, J. C. & Ackerman, S. L. Mutation of a U2 snRNA gene causes global disruption of alternative splicing and neurodegeneration. Cell 148, 296–308 (2012). 10.1016/j.cell.2011.11.057

55 Groschel, M. I., Sayes, F., Simeone, R., Majlessi, L. & Brosch, R. ESX secretion systems: mycobacterial evolution to counter host immunity. Nat Rev Microbiol 14, 677–691 (2016). 10.1038/nrmicro.2016.131

56 Stoop, E. J., Bitter, W. & van der Sar, A. M. Tubercle bacilli rely on a type VII army for pathogenicity. Trends Microbiol 20, 477–484 (2012). 10.1016/j.tim.2012.07.001

57 Vergne, I. et al. Mycobacterium tuberculosis phagosome maturation arrest: mycobacterial phosphatidylinositol analog phosphatidylinositol mannoside stimulates early endosomal fusion. Mol Biol Cell 15, 751–760 (2004). 10.1091/mbc.e03-05-0307

58 Del Rosario, R. C. H. et al. Histone acetylome-wide associations in immune cells from individuals with active Mycobacterium tuberculosis infection. Nat Microbiol 7, 312–326 (2022). 10.1038/s41564-021-01049-w

59 Denisenko, E. et al. Transcriptionally induced enhancers in the macrophage immune response to Mycobacterium tuberculosis infection. BMC Genomics 20, 71 (2019). 10.1186/s12864-019-5450-6

60 Louadi, Z. et al. DIGGER: exploring the functional role of alternative splicing in protein interactions. Nucleic Acids Res 49, D309–D318 (2021). 10.1093/nar/gkaa768

61 Bradley, R. K. & Anczukow, O. RNA splicing dysregulation and the hallmarks of cancer. Nat Rev Cancer (2023). 10.1038/s41568-022-00541-7

62 Boudreault, S. et al. Reovirus mu2 protein modulates host cell alternative splicing by reducing protein levels of U5 snRNP core components. Nucleic Acids Res 50, 5263–5281 (2022). 10.1093/nar/gkac272

63 De Maio, F. A. et al. The Dengue Virus NS5 Protein Intrudes in the Cellular Spliceosome and Modulates Splicing. PLoS Pathog 12, e1005841 (2016). 10.1371/journal.ppat.1005841

64 Almstead, L. L. & Sarnow, P. Inhibition of U snRNP assembly by a virus-encoded proteinase. Genes Dev 21, 1086–1097 (2007). 10.1101/gad.1535607

65 Corre, M. et al. Alternative splicing induced by bacterial pore-forming toxins sharpens CIRBP-mediated cell response to Listeria infection. Nucleic Acids Res 51, 12459–12475 (2023). 10.1093/nar/gkad1033

## Supplementary References

1 Liu, J. L. & Gall, J. G. U bodies are cytoplasmic structures that contain uridine-rich small nuclear ribonucleoproteins and associate with P bodies. Proc Natl Acad Sci U S A 104, 11655–11659 (2007). 10.1073/pnas.0704977104

2 Bolger, A. M., Lohse, M. & Usadel, B. Trimmomatic: a flexible trimmer for Illumina sequence data. Bioinformatics 30, 2114–2120 (2014). 10.1093/bioinformatics/btu170

68. Andrews, S. (2017).

4 Ewels, P., Magnusson, M., Lundin, S. & Kaller, M. MultiQC: summarize analysis results for multiple tools and samples in a single report. Bioinformatics 32, 3047–3048 (2016). 10.1093/bioinformatics/btw354

5 Patro, R., Duggal, G., Love, M. I., Irizarry, R. A. & Kingsford, C. Salmon provides fast and bias-aware quantification of transcript expression. Nat Methods 14, 417–419 (2017). 10.1038/nmeth.4197

6 Frankish, A. et al. GENCODE reference annotation for the human and mouse genomes. Nucleic Acids Res 47, D766–D773 (2019). 10.1093/nar/gky955

7 Love, M. I. et al. Tximeta: Reference sequence checksums for provenance identification in RNA-seq. PLoS Comput Biol 16, e1007664 (2020). 10.1371/journal.pcbi.1007664

8 Martin Morgan, V. O., Jim Hester, Hervé Pagès. SummarizedExperiment: SummarizedExperiment container. R package version 1.28.0 (2022). 10.18129/B9.bioc.SummarizedExperiment

9 Zhu, A., Srivastava, A., Ibrahim, J. G., Patro, R. & Love, M. I. Nonparametric expression analysis using inferential replicate counts. Nucleic Acids Res 47, e105 (2019). 10.1093/nar/gkz622

10 Vitting-Seerup, K. & Sandelin, A. IsoformSwitchAnalyzeR: analysis of changes in genome-wide patterns of alternative splicing and its functional consequences. Bioinformatics 35, 4469–4471 (2019). 10.1093/bioinformatics/btz247

11 Kolde, R. Pheatmap: pretty heatmaps. R package version v.1.0.12 (2019).

12 Chen, E. Y. et al. Enrichr: interactive and collaborative HTML5 gene list enrichment analysis tool. BMC Bioinformatics 14, 128 (2013). 10.1186/1471-2105-14-128

13 Ashburner, M. et al. Gene ontology: tool for the unification of biology. The Gene Ontology Consortium. Nat Genet 25, 25–29 (2000). 10.1038/75556

14 Wickham, H. ggplot2: Elegant Graphics for Data Analysis. Springer-Verlag New York (2016).

15 Team, R. C. R: A Language and Environment for Statistical Computing. (2022).

16 van Kessel, J. C. & Hatfull, G. F. Recombineering in Mycobacterium tuberculosis. Nat Methods 4, 147–152 (2007). 10.1038/nmeth996

17 Xu, C. et al. Structural insights into Gemin5-guided selection of pre-snRNAs for snRNP assembly. Genes Dev 30, 2376–2390 (2016). 10.1101/gad.288340.116

